# Informed Data-Independent Acquisition Enables Targeted Quantification of Key Regulatory Proteins in Cell Fate Decisions at Single-Cell Resolution

**DOI:** 10.1101/2025.05.30.656945

**Authors:** Jakob Woessmann, Valdemaras Petrosius, Sofie Ulfbeck Schovsbo, Benjamin Furtwängler, Tabiwang N. Arrey, Jeff Op de Beeck, Rahul Shanbhag, Nicholas Festy Glindtvad, Ajuna Azad, Eugen Damoc, Lars Velten, Bo T. Porse, Erwin M. Schoof

## Abstract

Low-abundance regulatory proteins, including transcription factors (TFs), remain largely inaccessible to direct quantification in single cells despite their central roles in cellular state transitions. Although single-cell proteomics by mass spectrometry (scp-MS) enables broad proteome profiling, current approaches often lack the sensitivity required to quantify these regulators at the protein level robustly. Here, we present informed data-independent acquisition (iDIA), a cross-instrument MS acquisition framework combining sensitive targeted measurements with global proteome profiling of the same cell. Thereby, iDIA significantly improves the sensitivity for predefined regulatory proteins, while preserving global proteome coverage. Applied to hematopoietic stem and progenitor cells, iDIA quantified 12 lineage-associated TFs, including GATA1 and SPI1, alongside the global proteome from single cells. Integrated analysis reconstructed the differentiation hierarchy and revealed protein-level states of early granulocytic-monocytic lineage priming and coordinated changes between erythroid TF abundance and cell-cycle progression. Thus, iDIA opens scp-MS to the regulatory architecture of cell state transitions.

## Introduction

Cellular differentiation is orchestrated by regulatory proteins such as transcription factors (TFs), chromatin modifiers, and signalling effectors, whose dosage and expression timing determine cell fate by regulating lineage-specific protein expression. These regulatory cell-state transitions have been described at single-cell resolution by single-cell RNA sequencing (scRNA-seq). However, mRNA abundance is only a proxy for the functional protein abundances that ultimately drive cell-fate decisions and we have previously shown weak mRNA-to-protein correlations in blood cells, especially in the most primitive stem cell compartment^1^. Direct quantification of the endogenous functional regulatory proteins throughout hierarchical cellular differentiation has so far been largely inaccessible at single-cell resolution.

Recent advances in single-cell proteomics by mass spectrometry (scp-MS) have substantially increased proteome depth and throughput. These gains, driven by improvements in sample preparation, chromatography, and mass spectrometry, have been accompanied by widespread adoption of data independent acquisition (DIA)^2–4^. Despite this progress, reproducible quantification of low-abundance proteins, including TFs, across a large number of heterogeneous cells remains challenging. As a result, protein-level TF dynamics in systems such as hematopoietic stem cell differentiation have continued to rely on bulk measurements averaging across millions of cells, masking the underlying cellular heterogeneity that drives lineage commitment^5^.

The complement to global DIA based scp-MS are sensitivity targeted-MS approaches such as parallel reaction monitoring (PRM). PRM has the potential to provide the required sensitivity to quantify TFs within single-cells, but its application in scp-MS remains limited and lacks global proteome context for further cell-state characterization^6,7^. Conversely, DIA enables global proteome coverage but still lacks sensitivity for robust low abundant TF quantification across large datasets.

Hybrid acquisition strategies that interleave PRM and DIA have proven powerful for bulk proteomics^8^, but their application to scp-MS is constrained by (1) dependencies on MS-instrument-specific implementations, (2) specific needs to accommodate long ion injection times (IIT) and (3) compatibility with fast chromatographic separations characteristic for scp-MS^2–4,9^. A hybrid acquisition strategy that is sensitive enough for single-cell application, broadly deployable across instruments and DIA search engines, and operable within standard LC/MS workflows has so far been lacking.

Here we introduce informed-DIA (iDIA), a generalizable acquisition framework that combines targeted and global proteome measurements in a single scp-MS experiment. iDIA methods are established through an open R-based web tool, removing the need of vendor application programming interfaces (APIs), allowing its application across Orbitrap (OT) platforms while remaining compatible with DIA search engines. Mechanistically, iDIA integrates targeted-MS2 scans into a DIA framework making it compatible with long IITs and fast chromatographic gradients.

We demonstrate that iDIA significantly reduces the limit of quantification (LOQ) for targeted peptides while maintaining a biologically interpretable global proteome. In combination with stable isotope-labelled (SIL) peptides, iDIA further improves quantitative robustness, enabling continuous quantification of low-abundant proteins across cellular differentiation. Applying the acquisition framework to study TF expression across the mouse hematopoietic stem and progenitor cell (HSPC) compartment, selected for its well-described TF expression pattern, iDIA enabled the characterization of expression dynamics of 12 TFs across HSPCs. By integrating the global proteome, the targeted TF panel, and antibody-based cell surface markers into a combined embedding, we resolve the canonical hematopoietic hierarchy. Critically, protein-level TF profiles separate early monocytic– and granulocytic-primed granulocyte-monocyte progenitor (GMP) subpopulations, while also revealing a coordinated coupling between erythroid-lineage TF abundance and cell-cycle dynamics.

## Results

### iDIA enables sensitive targeted quantification while retaining global proteome coverage

Regulatory proteins such as TFs are largely low-abundant and therefore only a small number can be quantified in low-input and single cell proteomics (**Figure 1A**). To address this limitation, we developed informed-DIA (iDIA), a platform-independent acquisition strategy that combines targeted and global proteome measurements within a single experiment for scp-MS (**Figure 1B**). In iDIA, targeted scans are not appended to the DIA acquisition method as previously reported^8^. iDIA instead replaces specific DIA MS2 isolation windows with PRM or wide-window PRM (wwPRM) scans^10^ (PRM scans including both endogenous and SIL peptide) (**Supplementary Figure 1 A, B**). To retain quantitative accuracy for precursors identified in DIA scans, MS1 scans are kept at a constant delta-time, and global proteome quantification is performed on MS1 level. The number of DIA MS2 windows that are replaced depends on the number of points-per-peak and the scheduled retention-time window for target peptides that the user aims for and can be adjusted in an experiment-specific manner. Furthermore, iDIA allows for customization of the targeted proteome acquisition with regards to number of targets, IIT, and number of DIA MS2 window replacements.

**Figure 1.**
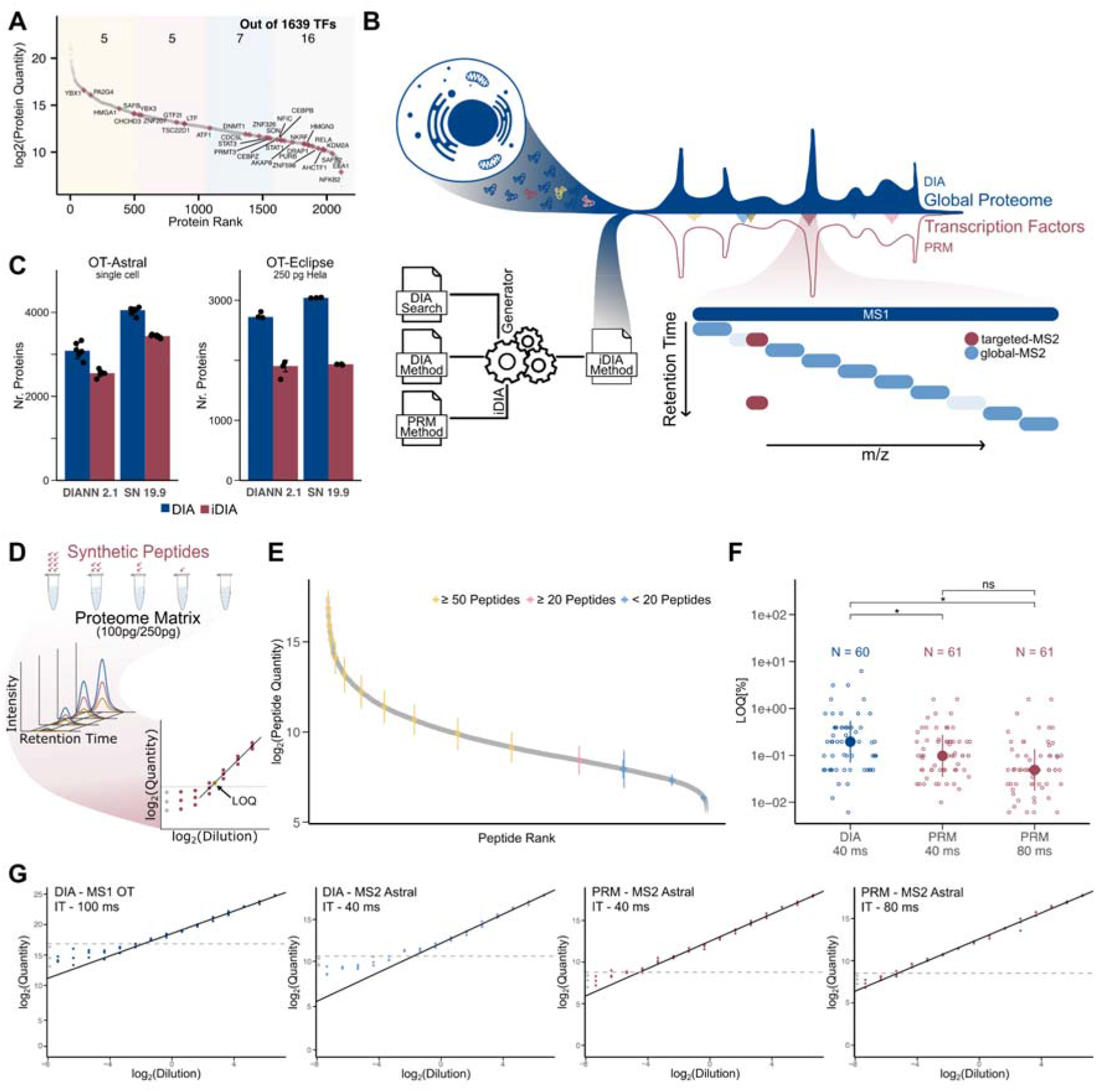
iDIA combines global and targeted proteomics at single cell resolution. (**A**) Ranked protein expression of 50 pg hela digest (n = 3). TFs identified which overlap with Lambert et al 2018^26^ highlighted. (**B**) Schematic of the iDIA Generator application and its generation of an iDIA method. Example of MS acquisition schema during iDIA acquisition of a single cell with the quantification of TFs and global proteome. (**C**) Comparison of the number of identified proteins by standard DIA (blue) and iDIA (red) with DIANN 2.1 and SN 19.9. Single HEK-293T cells acquired on a Orbitrap Astral Zoom, 9 theoretical peptides are acquired with targeted-proteomics replacing two out of 30 DIA windows per DIA cycle. 250 pg HeLa digest acquired on a Orbitrap Eclipse, 5 theoretical peptides are acquired with targeted-proteomics replacing 6 out of 6 DIA windows per DIA cycle. (**D**) Experimental setup to compare DIA and PRM sensitivity by determining the LOQ of synthetic peptides in a stable proteome matrix. (**E-G**) Quantitative comparison of PRM and DIA acquisitions on a Orbitrap Astral Zoom using a dilution curve of 61 synthetic peptides over 16 points in a 100 pg yeast and *e. coli* digest including blank, Median and MAD shown (**B**) Rank plot of mean peptide quantity of each peptide in the 100 pg yeast and E.coli digest with >66 % completeness across injections. Median and MAD of synthetic peptides identified at each dilution point. (**F**) LOQ of 61 synthetic peptides. Peptides were quantified via our gold standard DIA method based on MS1 peak area at 100 ms max IIT on the Orbitrap, PRM acquisition at 40 ms and 80 ms max IIT on the Astral Mass analyzer. T-test corrected by Benjamini & Hochberg (**G**) Example dilution curve of peptide EADDIVNWLK. Representative peptide closest to the median of all conditions. Summed raw peak area of top 3 y-ions displayed (3 replicates per dilution point). Peak area detected in negative control without synthetic peptides displayed in gray to the left. Mean + 2*sd around peak area without synthetic peptide shown as dashed line. *** p < 0.001, ** p < 0.01, * p < 0.05, ns p > 0.05

iDIA is set up directly in the method editor and can be implemented without the need for APIs or additional post-acquisition data processing. The iDIA Generator builds targeted mass list tables using a DIA method, a scheduled PRM method of SIL and/or endogenous peptides, and DIA search results of the sample of interest (or similar) as input. These targeted mass list tables are then loaded into the method editor, and the final data acquisition method is generated.

To demonstrate broad applicability, we deployed iDIA on a Thermo Scientific OT Astral Zoom and a Thermo Scientific OT Eclipse MS, and evaluated the compatibility of iDIA-generated raw files with two commonly used DIA search engines (DIA-NN ^11^ and Spectronaut). Single HEK-293T cells and 250 pg Pierce HeLa digest was acquired with standard DIA and iDIA containing 5 or 9 scheduled target peptides over the scheduled retention time of 0.5 or 0.7 min on both the OT Astral and the OT Eclipse. The resulting raw files were searched with Spectronaut 19.9 and DIA-NN 2.1 (**Figure 1C, Supplementary Figure 1 C,D**). On the OT Astral, 82.7 % (DIANN) and 84.7 % (SN) of the protein IDs were retained by iDIA, while 74 % and 72.3 % of the precursors remained. On the OT Eclipse, 70 % (DIANN) and 63.6 % (SN) of protein IDs were maintained (Precursors 58.4 % and 46.9 %). Thus, iDIA can be employed across MS instrumentation while preserving a DIA search engine-compatible global proteome.

### Targeted acquisition lowers the limit of quantification in low-input proteomics

Next, we assessed the quantitative sensitivity gain that targeted MS2 acquisitions included in the iDIA approach provided in low-input proteomics. We assessed the application of PRM acquisitions across two LC-MS setups in two sample matrices. Sensitivity of PRM and DIA acquisitions at various IITs were measured through dilution curves of synthetic peptides in a stable proteome (≤250 pg on column). Based on the quantitative linear range, we determined the LOQ of the synthetic peptides as a measure of sensitivity. We utilized our scp-MS optimized DIA acquisition methods as baseline to explore whether targeted MS2 scans could extend the LOQ in low-input proteomics (**Figure 1D**). To evaluate the broad applicability, we used two independent synthetic peptide pools acquired on an OT Eclipse and OT Astral Zoom. Experiments on the OT Eclipse were performed with 10 synthetic peptides and their equivalent SIL versions, diluted in a stable 250 pg HeLa digest. On the OT Astral Zoom 61 synthetic peptides were diluted in a 100 pg Yeast and E.coli digest. The synthetic peptides were diluted in 16 steps through the respective proteome matrix, covering the entire dynamic range of the global proteome. DIA quantification was performed on the MS1 level on both instruments due to the higher number of points per peak that could be reached. PRM acquisitions of different max IITs were kept at a stable points-per-peak to ensure consistent accuracy and precision, with results only impacted by max IIT (**Supplementary Figure 1E**).

We observed that PRM acquisitions reached significantly lower median LOQs compared to DIA across both platforms. On the OT Eclipse, targeted acquisitions improved the median LOQ by 1.3-fold at 118 ms and 2.5-fold at 246 ms IIT compared to DIA (**Supplementary Figure 1F**). On the OT Astral Zoom, median LOQ was improved median two-fold at 40 ms and four-fold at 80 ms compared to DIA (**Figure 1F, G**). On both instruments, longer IITs further decreased the LOQ. Overall, tailored targeted acquisition methods decreased the median LOQ of peptides in low-input proteomics across instruments.

### Wide-window PRM enables multiplexed and robust quantification in low-input samples

In order to improve the reproducibility of peptide quantification, the addition of SIL peptides in targeted MS acquisitions is common practice^12^. SIL peptides can be used to control for LC/MS and sample matrix related variations during acquisition related to sensitivity losses and sample injection across runs in scp-MS.

We evaluated the necessity of SIL peptides for quantification in low-input proteomics by assessing quantitative linearity of 10 synthetic peptides (**Supplementary Figure 1G**). Nine out of 10 peptides could be fitted with a linear standard curve when using a spiked-in SIL peptide for normalization, compared to five out of 10 peptides using label-free quantification alone (**Supplementary Figure 1H**). However, the acquisition of both SIL and endogenous peptides with individual PRM scans leads to a doubling of the required targeted MS2 scans and as a consequence, limits target multiplexing. Therefore, we explored the application of wwPRM in low-input proteomics.

The widening of the isolation window did not significantly impact the overall LOQ in low-input proteomics, while enabling simultaneous acquisition of SIL and endogenous peptides, reducing the MS duty cycle and allowing the incorporation of more targeted peptides. (**Supplementary Figure 2A, B**). We further observed reduced coefficients of variation (CV) for wwPRM acquisitions compared to DIA across the dilution series (**Supplementary Figure 2C**). Together, these results demonstrate that SIL peptides in combination with wwPRM enable multiplexed and robust quantification in low-input proteomics without compromising sensitivity.

### iDIA balances sensitivity and proteome coverage in low-input workflows

Having demonstrated the ability to combine global proteome and highly sensitive targeted proteome measurements in a single MS experiment using iDIA, we next evaluated the performance of iDIA in balancing targeted sensitivity with global proteome coverage. We assessed the applicability of iDIA based on three DIA methods in a 50 pg HeLa digest matrix spiked with 10, 20 or 34 SIL peptides acquired over 30-second scheduled wwPRMs on an OT Astral (**Supplementary Table 1**, **Supplementary Table 2**) (**Figure 2A**). Below we demonstrate the impact of (1) number of targeted peptides, (2) the IIT of targeted MS2 acquisition, (3) global proteome quantification and (4) targeted peptide points-per-peak on an iDIA acquisition.

**Figure 2.**
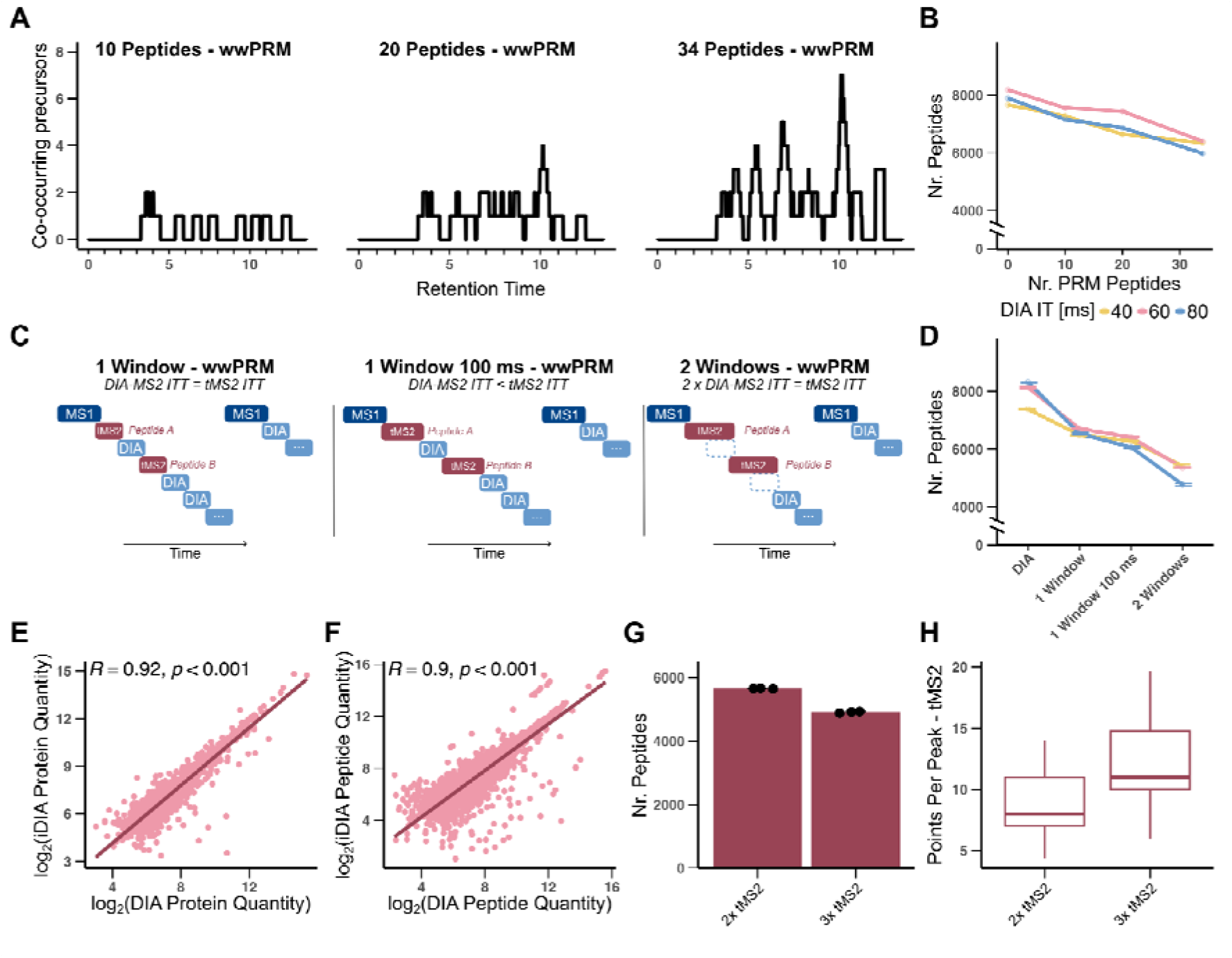
iDIA enables balance between global and targeted proteome coverage. iDIA performance evaluation in 50 pg HeLa digest 34 SIL peptides were spiked into 50 pg HeLa digest and acquired with iDIA methods based on three DIA methods displayed in Table 1. (**A**) Number of cooccurring precursors which are acquired in wwPRM scans during iDIA method. (**B**) Number of peptides identified in triplicate acquisition of iDIA method quantifying 10, 20 or 34 peptides with wwPRM scans compared to DIA. (**C**) Schematic of iDIA acquisition options. Replacement of a DIA window with a wwPRM scan, replacement and extension of maxIIT of a DIA window with a wwPRM scan or replacement of two DIA windows by one wwPRM scan. (**D**) Number of peptides identified in triplicate acquisition of iDIA methods shown in C. (**E, F**) Pearson Correlation of peptide and protein level quantification of peptides identified in DIA search during iDIA acquisition. Correlation between example DIA acquisition and iDIA acquisition with a 60 ms max IIT and 20 window DIA method. (**G, H**) iDIA acquisition with 2 or 3 DIA MS2 windows replaced per MS2 scan cycle. (**G**) Number of peptides identified by SN 19.9 (**H**) Points-per-peak of 34 peptides acquired in wwPRM scans during iDIA acquisition.

The increasing number of target peptides in the iDIA method led to an increased loss of peptide and protein identifications in the global proteome compared to standard DIA (**Figure 2B, Supplementary Figure 3A**). While 95 % of the global proteome was retained when 10 targeted peptides were included, a maximum of 24.2 % of the global proteome was lost when acquiring 34 peptides in wwPRM during iDIA acquisition at 80 ms max IIT with the lowest number of MS2-DIA isolation windows.

Next, we assessed the impact of increasing IIT of target-acquisitions on the global proteome coverage. To allow full flexibility, the online iDIA generator enables three approaches of DIA-MS2 window replacement by targeted-MS2 scans (**Figure 2C**). DIA windows can either 1) be replaced by a wwPRM scan (same IIT of wwPRM and DIA MS2), 2) the wwPRM scan window can be extended (longer IIT of wwPRM than DIA MS2) or 3) the wwPRM scan can replace two consecutive DIA windows (wwPRM scans have twice the IIT of DIA scans in MS2). The extension of the wwPRM IIT to 100 ms (Option 2) reduced the number of identified global peptides on average by 4.9 % compared to standard iDIA (Option 1). Replacing two consecutive DIA MS2 windows (Option 3) further decreased the global peptide coverage by 21.2 % (**Figure 2D, Supplementary Figure 3B**). The number of MS1 points-per-peak was kept stable in all iDIA and DIA methods, ensuring robust MS1-based quantification (**Supplementary Figure 3C**). The lower the number of DIA MS2 windows remaining in the DIA methods, the more peptides were lost in the global proteome, highlighting the inherent balance between DIA MS2 isolation windows and DIA IIT.

When comparing the quantitative performance of iDIA (Option 2) with its corresponding standard DIA acquisition, we highlight that the global proteomes correlate significantly (Pearson correlation of 0.92 at the protein level and 0.9 at the peptide level), with only a minor reduction compared to DIA acquisitions (Pearson correlation of 0.96) (**Figure 2E,F, Supplementary Figure 3D,E**).

Finally, iDIA allows for custom specification of points-per-peak for the targeted-proteome. This is achieved by replacing a larger proportion of DIA-MS2 scans with targeted MS2 scans (in our example we replaced 2 and 3 out of 20 DIA scans per MS2 scan-cycle for each target peptide). We could observe a decrease of peptide IDs by 13.1 % from a mean of 5657 to 4914 identified global peptides (**Figure 3 G**). Meanwhile, the median number of points per peak of the targeted proteome increased from 8 to 11 (**Figure 3 H**). These increases in wwPRM windows lead to a decrease (not significant) in median LOQ (**Supplementary Figure 3 F**). Overall iDIA provides a highly customizable acquisition strategy that can balance the global– and targeted proteome to individual experimental requirements.

**Figure 3.**
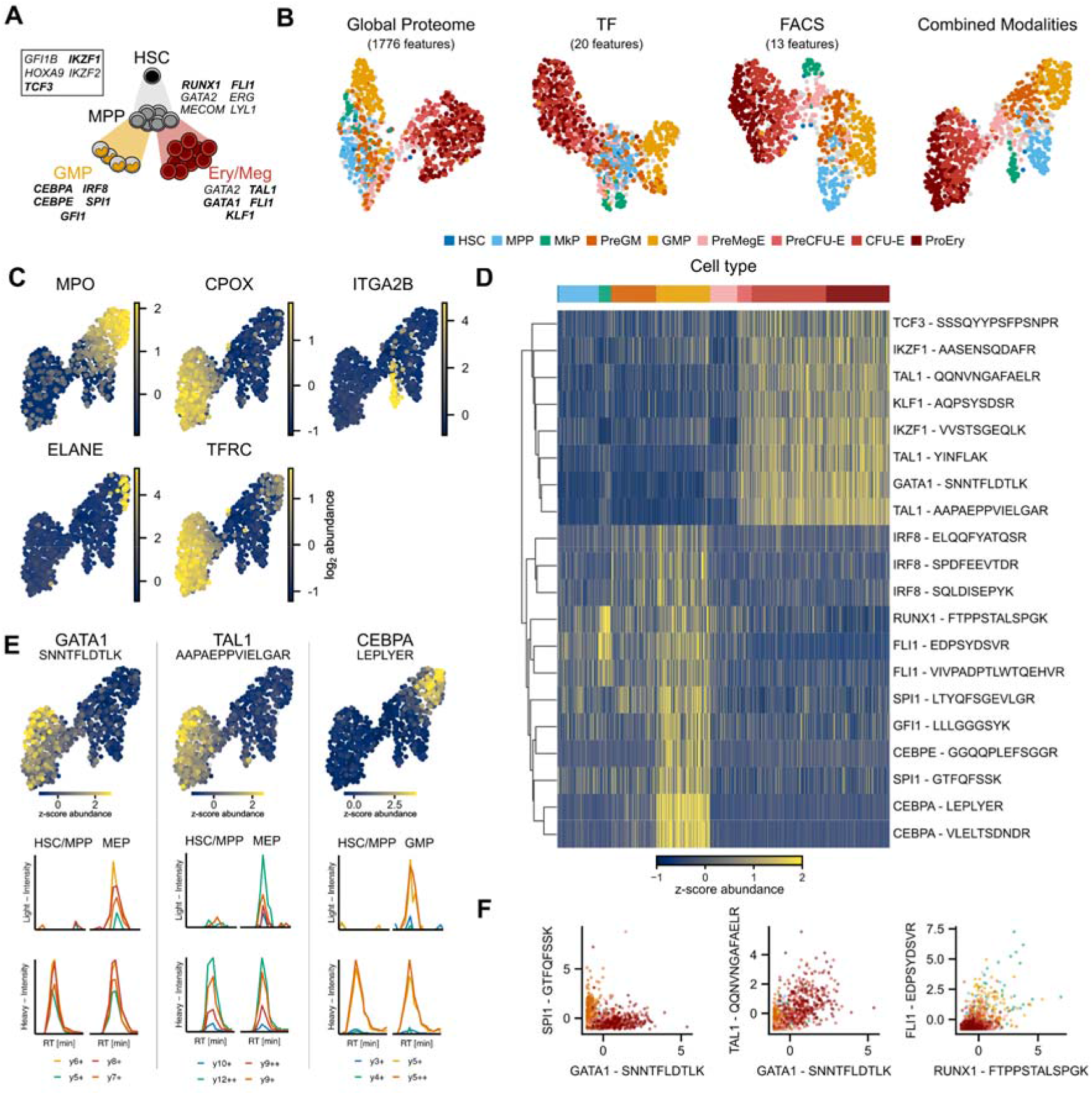
iDIA quantifies HSPC heterogeneity and lineage specific TFs at protein level in single HSPCs. **(A)** cKIT+Lin– HSPC compartment with lineage specific TFs targeted with iDIA. TFs that were quantified across single cells highlighted in bold. **(B)** UMAP of global proteome, TFs, FACS cell surface markers and combined embedding using WNN acquired from the same HSPC. Grey cells without cell type annotation. See Supplementary Figure 7C for extended annotation. **(C)** UMAPs of lineage specific log2 protein abundance of proteins quantified in global proteome **(D)** Cell-type specific z-scored TF abundance at peptide level. Hierarchical clustering of TFs. **(E)** UMAP of lineage specific z-scored TF abundance. Example raw XIC quantification of TF peptide in HSC/MPP and erythroid-megakaryocyte progenitors or GMP population. Bottom SIL reference peptide intensity top endogenous peptide intensity in single cell. **(F)** Correlation between lineage specific z-scored TF abundance across cell-types.

### iDIA enables robust quantification of transcription factors in single mouse hematopoietic stem and progenitor cells

The hematopoietic stem and progenitor cell (HSPC) hierarchy constitutes a continuous differentiation system where cell lineage fate decisions are (in part) regulated by the actions of TFs. Quantifying TFs directly at the single-cell protein level imposes a particularly stringent sensitivity requirement on scp-MS due to their overall low abundance, as shown in **Figure 1A**. In addition, primitive HSPCs are among the smallest mammalian cells^13^. We therefore selected the mouse HSPC compartment as a deliberately stringent test case for iDIA, reasoning that a framework capable of resolving TF dynamics in this regime would be broadly deployable across biological systems in which low-abundance regulators determine cell state.

To test this hypothesis, we used flow cytometry to sort cKIT+Lin– mouse bone marrow HSPCs (**Supplementary Figure 4**). cKIT+Lin– cells contain hematopoietic stem cells (HSCs), which differentiate into multipotent progenitors (MPP) and further bifurcate into lineage-committed progenitors including granulocyte-monocyte progenitors (GMPs) and a variety of erythroid-megakaryocyte progenitors. To characterize the HSPC compartment, we developed a transcription factor panel of 34 peptides covering 19 TFs based on previously acquired data and literature^5,14,15^ (**Supplementary Table 2, Figure 3A**). The panel was integrated into an iDIA acquisition method using wwPRM scans in combination with SIL peptides.

First, we assessed the ability of iDIA to not only retain global proteome coverage, but also its ability to deliver precise TF quantification across varying cell inputs. In single cKIT+Lin– cells, we quantified on average 1092 proteins per cell using standard DIA and 817 proteins with the iDIA method, demonstrating retention of global proteome coverage in primary cells (n = 12) (**Supplementary Figure 5 A, B**). To evaluate quantitative performance across input amounts, we analyzed 0, 1, 5 and 10 pooled cKIT+Lin– cells with iDIA. Global proteome detection increased with cell input as expected (**Supplementary Figure 5 D, E**), and 24 peptides across 14 TFs were identified within the targeted proteome. The TF abundances broadly followed the input cell number (**Supplementary Figure 5 F**). Due to the heterogeneous nature of the HSPC compartment, the quantified TFs in pooled 5 and 10 cell iDIA acquisitions are not expected to scale linearly. Across input amounts, we obtained median CVs of 4.9% for the SIL peptide spike-in, highlighting the robustness of the SIL peptide quantification of TFs in single cells (**Supplementary Figure 5 C**). Together, these benchmarks allowed us to conclude that iDIA retains global proteome coverage and delivers precise targeted TF quantification in low-input HSPC samples.

To showcase the applicability of iDIA to individual cells isolated from the HSPC compartment, we next single-cell sorted cKIT+Lin– HSPCs while retaining the FACS information from an 11-plex cell surface marker panel which we used to delineate cell populations by conventional FACS gating (Pronk et al. 2007^16^)(**Supplementary Table 3, Supplementary Figure 6-7A**). We subsequently used iDIA to acquire an scp-MS dataset of 905 cells, resulting in a global proteome containing 1776 quantified proteins (719 median per cell) alongside the simultaneous quantification of 12 TFs with 20 peptides in single HSPCs with a TF (targeted-proteome) data missingness of 3.5 % (median). Together, these results demonstrate that iDIA enables simultaneous global proteome profiling and robust low-missingness quantification of key TFs in single primary mouse HSPCs, thereby providing a framework to further explore the quantified TF expression.

### iDIA enables multi-modal integration to resolve the HSPC hierarchy

We first generated low-dimensional UMAP embeddings of the global proteome, the targeted TFs and the flow-cytometry cell surface markers (**Figure 3B**). Each recapitulated the hematopoietic hierarchy and separated HSC/MPPs, erythroid-megakaryocyte progenitors and GMPs. Additionally, each modality highlighted TF expression patterns with subtle nuances distinct to each modality (**Supplementary Figure 8-10**). To obtain the most objective molecular representation of the hierarchy, we reasoned that a combined embedding of the global proteome, the TF proteome and the FACS markers using weighted nearest neighbor (WNN)^17^ would harmonize TF expression patterns and allow for an integrated analysis (**Figure 3B, Supplementary Figure 7B, 11, 12**). The embedding of the combined modalities recapitulated HSPC differentiation and displayed a clear differentiation hierarchy of MPPs differentiating in the respective progenitors that lead into separate erythrocytic-megakaryocytic and granulocytic-monocytic branches. Notably, the additional separation of MkP progenitors could be identified which is also present in the TF and FACS embedding. The global proteome also recapitulated known lineage specificity and cell-type specific protein markers for GMPs (MPO, ELANE) and erythrocytic-megakaryocytic progenitors (CPOX, TFRC, ITAG2B) (**Figure 3C, Supplementary Figure 13**). Furthermore, we could identify cell-type and –lineage specific expression of all quantified TFs for all peptides. While we observed common erythrocytic cell types being defined by TFs such as TAL1, GATA1, TCF3 and KLF1, GMPs displayed characteristic expressions of SPI1, CEPBA, CEBPE and IRF8 (**Figure 3D-E, Supplementary Figure 14**). Notably, lineage-affiliated TFs such as SPI1 and GATA1 showed clear anticorrelation between the erythroid-megakaryocyte and GMP branch. TFs specific for the same lineage, such as TAL1 and GATA1, displayed correlation in the erythroid-megakaryocyte progenitor lineage, and we could identify TF-pairs which only show combined expression in specific cell-types such as FLI1 and RUNX1 in MkPs and otherwise were not or lowly expressed in combination (**Figure 3F**).

### Protein-level transcription factor dynamics resolve lineage commitment in hematopoiesis

We further investigated the lineage specificity of TFs by calculating the pseudotime for the granulocytic-monocytic and the erythrocytic differentiation branch (**Figure 4A**). First, we explored MkP specific TF expression which is uniquely different to the further erythrocytic lineage. We observed a significantly different expression of TAL1, FLI1 and RUNX1 in MkPs vs. other progenitor populations with a notable significant downregulation of IKZF1 (**Figure 4B**). We next assessed the granulocytic-monocytic and erythrocytic branch and found that different peptides quantifying the same TF displayed the same quantitative trend across pseudotime, highlighting the strong quantitative ability of iDIA. We partially highlight clear lineage-specific expression of all TFs except FLI1 and IKZF1 (**Figure 4C**). While FLI1 displayed a downregulation over pseudotime in the erythroid branch and a strong upregulation in the granulocytic-monocytic branch, IKZF1 expression increased in both branches. IKZF1 was stronger expressed in the erythroid branch while showing an earlier but weaker upregulation in the granulocytic-monocytic branch. The granulocytic-monocytic branch displayed an overall upregulation of CEBPA followed by a strong CEBPE and GFI1 expression. Notably, IRF8 displayed an initial upregulation followed by downregulation across pseudotime.

**Figure 4.**
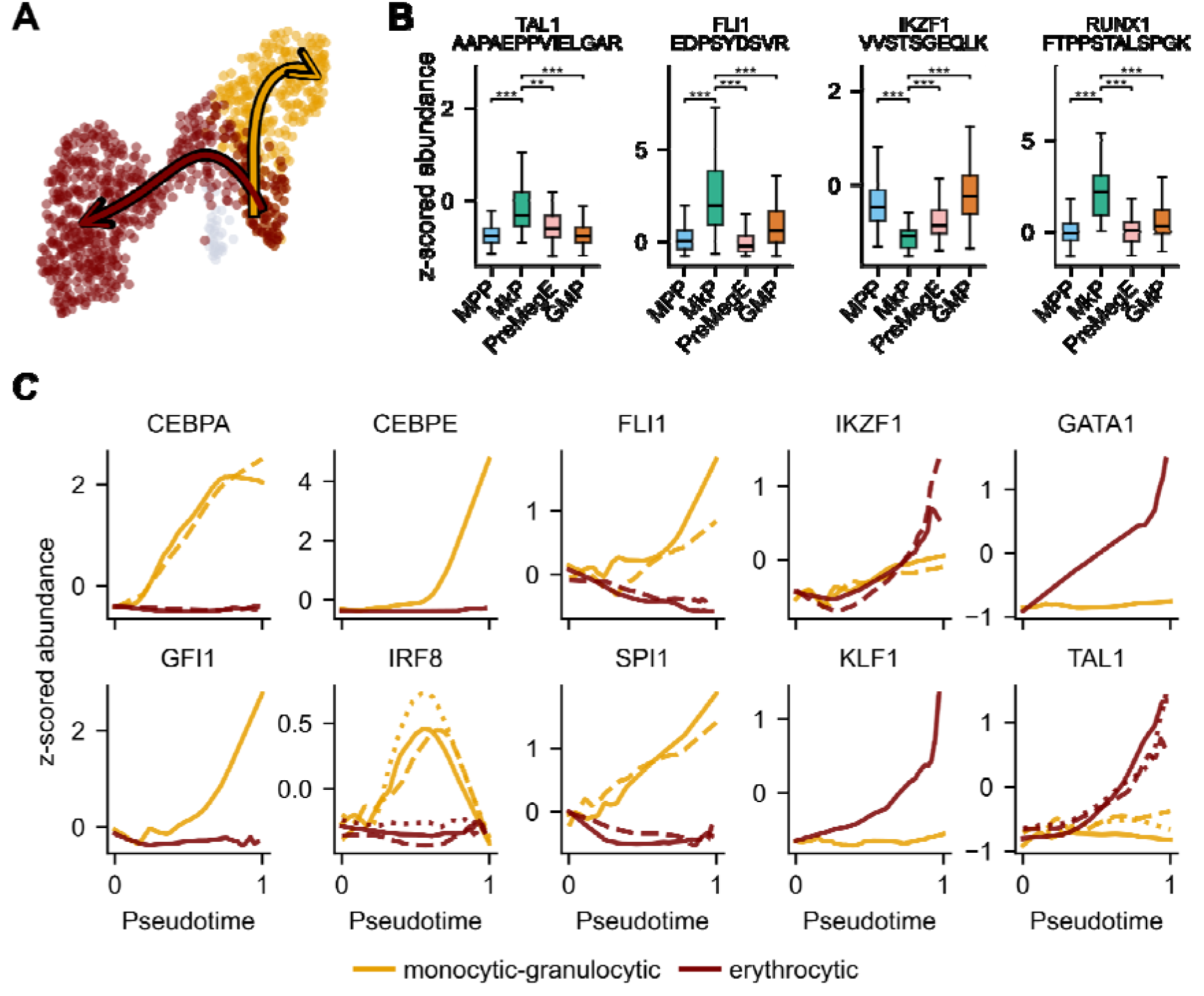
iDIA deciphers lineage specific TF expression along pseudotime in HSPCs. **(A)** erythrocytic and monocytic-granulocytic pseudotime from primitive MPP/HSC. **(B)** Significantly up or downregulated TF in MkP cells compared to MPP, PreMegE and GMP by Mann-Whitney-U-test corrected for multiple testing by fdr. *** p < 0.001, ** p < 0.01, * p < 0.05, ns p > 0.05 **(C)** Z-scored abundance of TFs over pseudotime along erythrocytic and monocytic-granulocytic differentiation branches. Dashed lines show multiple peptides quantified of the same TF across single cells see Supplementary Table 2.

### TF expression captures early separation of early monocytic and granulocytic differentiation in GMPs

We next focused on the GMP population (GMP leiden cluster) and clustered cells based on their expression of the GMP specific TFs CEBPA, CEBPE, IRF8, SPI1 and GFI1. TF expression clearly separated GMPs into what we termed bi-lineage GMPs (limited expression of GFI1, CEBPE and IRF8), granulocytic progenitors (expression of GFI1 and CEBPE) and monocytic progenitors (expression of IRF8) (**Figure 5A**). These monocytic– and granulocytic primed progenitors displayed clear separation in UMAP space in the combined modalities and at TF level, however, they could not be directly separated in the individual global proteome and FACS surface-marker embeddings (**Figure 5B, Supplementary Figure 8-12**). Monocytic– and granulocytic-primed progenitor cells displayed distinct expressions of either IRF8 or GFI1 and CEBPE while being overall CEBPA positive (**Figure 5C, Supplementary Figure 14**). Turning to the global proteome of these two cell clusters we could identify a distinct monocytic (e.g. F13A1) and granulocytic (e.g. ELANE, ANXA1)^18–20^ protein expression profile between the two groups (**Figure 5D**). Taken together we could highlight a separation of GMPs into monocytic, granulocytic and bi-linage progenitors facilitated by the iDIA-derived TF expression patterns.

**Figure 5.**
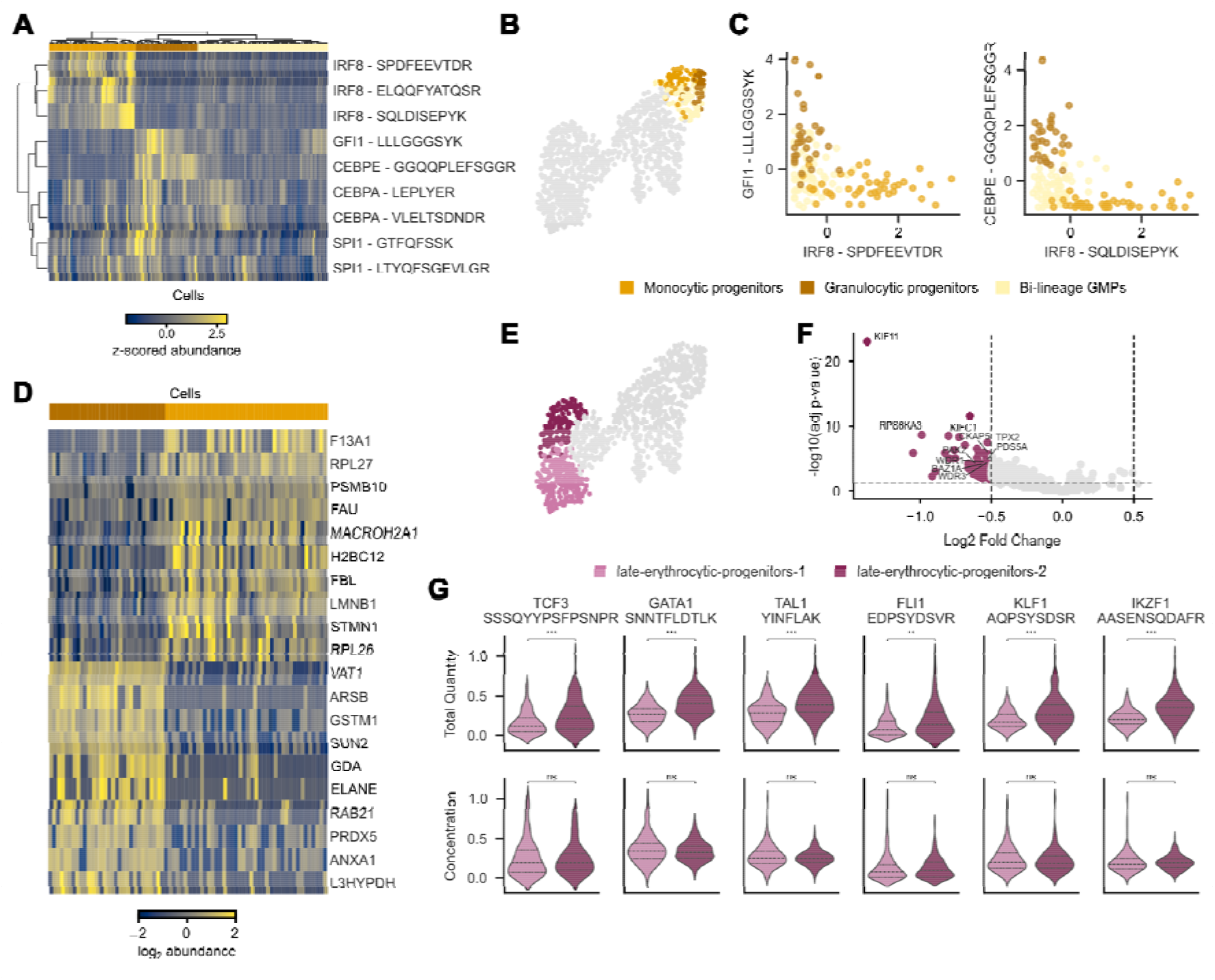
TF expression differentiates monocytic and granulocytic as well as late-erythrocytic progenitors. **(A)** GMP specific z-scored TF abundance clustered at peptide and cell level. Three main cell clusters within the leiden GMP cluster were annotated as Monocytic-, Granulocytic– and Bi-lineage-Granulocytic-Monocytic– Progenitors **(B)** UMAP of combined modalities highlighting GMP subclusters **(C)** Subcluster specific z-scored TF abundance profile highlighting distinct Monocytic-and Granulocytic-Progenitors. **(D)** Differentially express proteins of the global proteome between Monocytic-and Granulocytic-Progenitors (10 highest fold-changes between groups displayed) **(E)** UMAP of combined modalities highlighting leiden subcluster of Erythrocytic differentiation branch **(F)** Differentially expressed proteins between erythrocytic subclusters. Significantly upregulated proteins involved in mitotic spindle formation and cell cycle regulation highlighted. **(G)** TF abundance of erythroid lineage TFs between subclusters. Total quantity of TF in cells and TF concentration (total quantity/total-MS-signal) displayed. Mann–Whitney-U-test corrected for multiple testing by fdr. *** p < 0.001, ** p < 0.01, * p < 0.05, ns p > 0.05

### Erythrocytic lineage TFs suggest cell-cycle dependent regulatory mechanisms

In the erythrocytic lineage branch we observed a separation of ProEry and CFU-E by their TF expression into high and low expression of erythroid lineage specific TFs and a separation of the cells into two leiden clusters (**Figure 5E, Supplementary Figure 7D and 12**). The two late erythrocytic progenitor clusters differentially expressed 59 proteins with proteins related to mitotic spindle formation and cell cycle regulation (**Figure 5F**). This suggested that late erythrocytic progenitors were separated by cellular proliferation. Notably we observed a significant increase in TF expression between the two clusters with higher total amounts of TF present in the cycling cells (**Figure 5G**). However, when normalizing the TF expression to the overall global proteome signal (i.e., the TF concentration within the overall proteome) we observed no significant difference between the two late erythrocytic progenitor clusters, suggesting a possible buffering of TF levels throughout the cell cycle.

## Discussion

In this study, we developed iDIA, a single-cell-optimized LC/MS acquisition framework that integrates targeted-peptide acquisitions into a DIA backbone to combine discovery proteomics with at least a four-fold increased sensitivity of targeted quantification within a single experiment. By replacing selected DIA-MS2 scans with targeted-MS2 scans rather than appending them, iDIA accommodates the long ion injection times within short chromatographic peak widths that are commonly used in scp-MS^4^. By operating through the standard method editor and generating raw files that are searchable by commonly used DIA search engines, it sidesteps instrument-specific APIs and allows for direct application across instruments. We highlighted this cross-platform deployment on the OT Astral, OT Astral Zoom and OT Eclipse. Together we demonstrated that iDIA is a generalizable MS-acquisition framework with novel global– and targeted MS2 scan integration tailored towards increased sensitivity and quantitative performance.

Applied to the mouse HSPC compartment, iDIA enabled the first direct, non-antibody-dependent protein-level quantification of lineage-defining TFs across single hematopoietic progenitors. Obtaining the global proteome, targeted TF panel, and surface-marker expression profile of each cell highlighted that each modality recapitulated the established HSPC differentiation hierarchy, while contributing with unique features that allowed a combined multi-modal analysis. Canonical antagonistic TF pairs such as GATA1/TAL1 versus SPI1/CEBPA/CEBPE, were resolved at the protein level along the erythroid-megakaryocytic and granulocytic-monocytic branches respectively, consistent with prior literature^5,15,21^. More notably, protein-level TF profiles separated GMPs into early monocytic– (IRF8-high), granulocytic– (GFI1/CEBPE-high), and bi-lineage progenitor states as previously inferred and validated from scRNA-seq lineage-tracing^15,19,22^ but, which had not been directly confirmed at the protein level. Notably, the hypothesis-driven TF panel enabled this observation while the discovery-driven global proteome expression of the same cells allowed for its confirmation, demonstrating that the targeted-TF layer added by iDIA unlocks regulatory cell-state information that remains invisible to global scp-MS readouts alone. In the erythroid branch, late-progenitor clusters distinguished by proliferation status showed elevated absolute TF abundance but stable TF-to-total-proteome concentration, suggesting that the quantified erythroid TFs are buffered against the total-proteome scaling that accompanies cell-cycle progression. Furthermore, a significant downregulation of IKZF1 in MkPs relative to all other progenitor compartments, consistent with the established repressive role of Ikaros in megakaryopoiesis was revealed^23^. Together, these findings extend our recent multimodal mapping approach of early human blood differentiation^1^ by integrating a targeted-TF layer that scp-MS has been lacking until now.

iDIA enables the direct targeted quantification of proteins with limited to no data-missingness at single-cell resolution with significantly increased sensitivity compared to global protein quantification. This is fundamentally distinct from the most widely used alternatives to study lineage regulation based on RNA-sequencing or affinity-based approaches^15,21^. Additionally, the ability of iDIA to quantify peptides based on SIL peptides will enable not only relative quantification as presented in this study but also allow to report copy numbers per cell based on absolute-quantified SIL peptides. Moreover, the iDIA framework is agnostic to target identity and its use will extend well beyond TFs in hematopoiesis. Specifically, we anticipate applications where sensitive protein measurements with low data missingness are needed, which can include, among others, the quantification of post-translational modifications or surface marker panels at single-cell resolution.

Having demonstrated that iDIA enables direct protein-level quantification of regulatory cell-state transitions in primary HSPCs, several limitations are worth noting. First, iDIA requires prior knowledge of the targets of interest. While this is a strength for hypothesis-driven studies in well-characterized systems, fully unbiased discovery of low-abundant proteins will continue to rely on complementary bulk reference experiments. Second, the targeted peptide panel size is currently bounded by the scheduling constraints of high-throughput LC-MS gradients. In our HSPC study, acquired at ∼85 samples-per-day, a maximum of 6 co-eluting precursors could be scheduled, comfortably accommodating 34 peptides covering 19 TFs. Several developments may address these limitations in future implementations. Adaptive retention time alignment could increase tolerance to chromatographic variability, enabling narrower precursor scheduling in large-scale experiments, and would concomitantly allow for increasing the targeted peptide panel size. In parallel, improvements in instrument sensitivity could enhance both targeted peptide detection and global proteome depth. Analysis throughput may be further increased through integration with multiplexed acquisition strategies, such as plexDIA^24^ or mDIA^25^.

Together, these results demonstrate that targeted and discovery proteomics can be effectively combined within a generalizable single-cell LC/MS acquisition and analysis strategy. By enabling direct reproducible quantification of low-abundance regulatory proteins alongside the global proteome, iDIA provides cross-MS-platform access to a molecular layer that has remained largely inaccessible to current scp-MS workflows. These developments will have broad implications for our ability to understand cell fate decisions across biology.

## Acknowledgements

We would like to thank the DTU Bioengineering Data Science Hub for support with online application setup and Zixiang Pan for critical input on the data processing. Work in the Porse lab was supported by grants from the Independent Research Foundation Denmark, The Novo Nordisk Foundation, The Danish Cancer Society and the Eva and Henry Frænkel Memorial Foundation. The project is part of the Danish Research Center for Precision Medicine in Blood Cancers funded by the Danish Cancer Society grant no. R223-A13071 and Greater Copenhagen Health Science Partners. Work in the Schoof lab was funded by the following grants: 1) reference number NNF21OC0071016 from the Novo Nordisk Foundation; 2) case no. 2067-00053B from the Independent Research Fund Denmark, 3) Lundbeck Foundation (R413-2022-869), 4) Velux Foundation (00053026), and 5) the DigitSTEM initiative with Bioneer. Jakob Woessmann acknowledges support from the Danish Ministry of Higher Education and Science for the Elite Research travel grant.

## Data availability

MS raw data and data matrices are available through panorama public (https://panoramaweb.org/iDIA.url) with the ProteomeXchange ID PXD079840.

## Code availability

All Code for the generation of the manuscript is available through GitHub as well as panorama public (https://panoramaweb.org/iDIA.url).

---

**Supplementary Figure 1.**
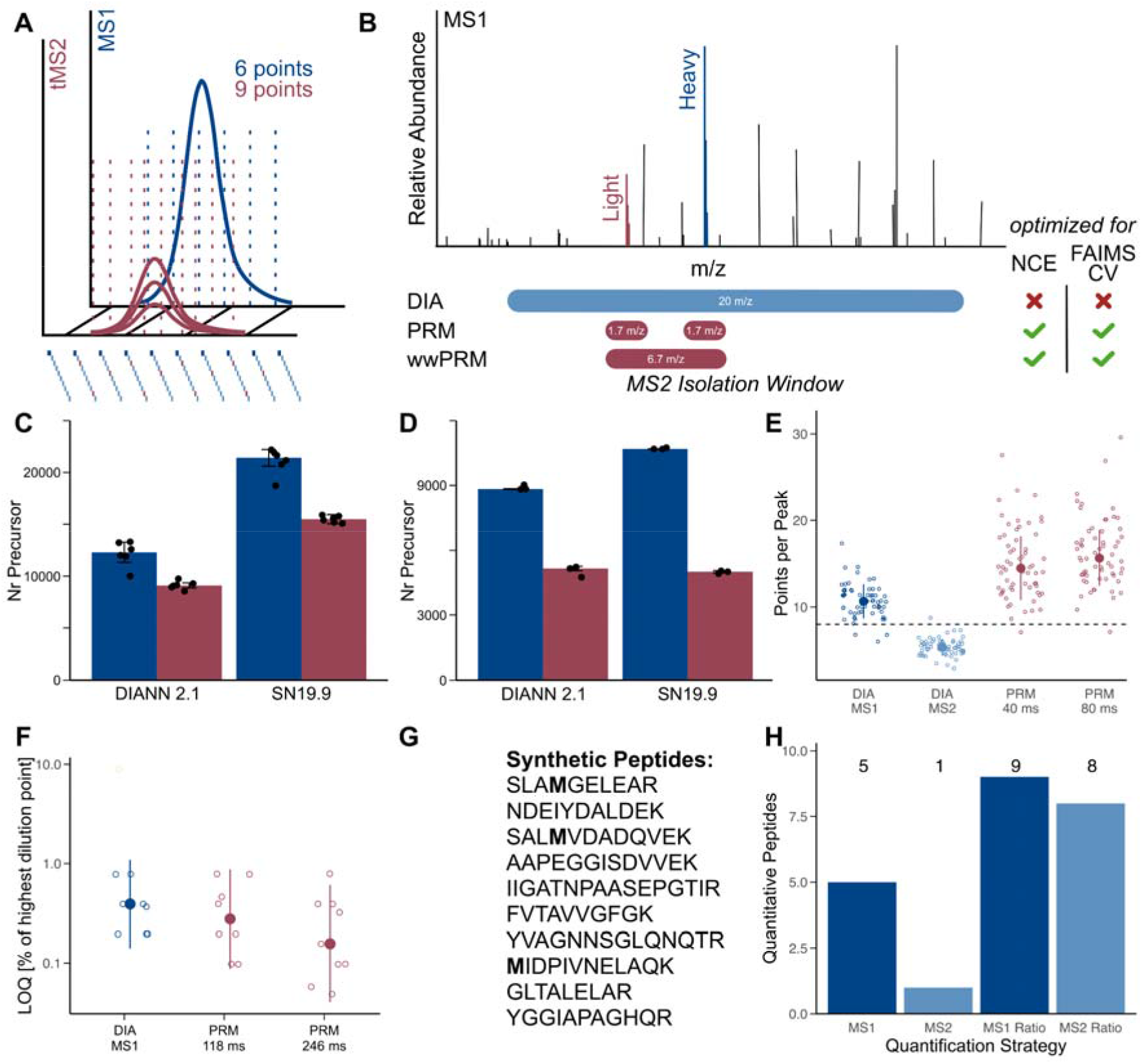
iDIA combines global and targeted proteomics at single cell resolution. (**A**) iDIA acquisition approach. DIA acquisitions (MS2) are performed in combination with targeted PRM/wwPRM scans (tMS2). tMS2s replace DIA scans to maximize points-per-peak for the tMS2 scans. DIA-quantification is performed at the MS1 level to maintain global proteome points-per-peak. (**B**) Theoretical MS1 spectrum containing a peptide that is present both as endogenous (light) and SIL (heavy) peptide. MS2 isolation windows of DIA, PRM and wwPRM acquisitions displayed. (**C-D**) Comparison of the number of identified precursors by standard DIA (Blue) and iDIA (Red) with DIANN 2.1 and SN 19.9. (**C**) Single HEK-293T cells acquired on a Orbitrap Astral Zoom, 9 theoretical peptides are acquired with targeted-proteomics replacing two out of 30 DIA windows per DIA cycle. (**D**) 250 pg HeLa digest acquired on a Orbitrap Eclipse, 5 theoretical peptides are acquired with targeted-proteomics replacing 6 out of 6 DIA windows per DIA cycle. (**E**) Points-per-peak of 61 peptides between integration boundaries in Skyline. PRM points-per-peak were kept comparable by means of MS2 dummy scans. The dashed line indicates 8 points per peak. (**F**) LOQ of 10 synthetic light peptides quantified by ratio to SIL peptides on the Orbitrap Eclipse. Peptides were quantified via our gold standard DIA method based on MS1 peak area at 246 ms max IIT, PRM acquisition at 118 ms and 246 ms max IIT. Median and MAD are shown for each condition. (**G**) List of 10 peptides synthesized as SIL and light randomly selected from the *Arabidopsis thaliana* proteome. (**H**) Number of peptides shown in G that decreased linearly in dilution curves and to which a standard curve could be fitted. Peptides quantified in DIA on an Orbitrap Eclipse. SIL peptide was kept at a stable concentration throughout the dilution curve. “Ratio” refers to quantification of light peptide in ratio to stable spike-in of SIL peptide.

**Supplementary Figure 2.**
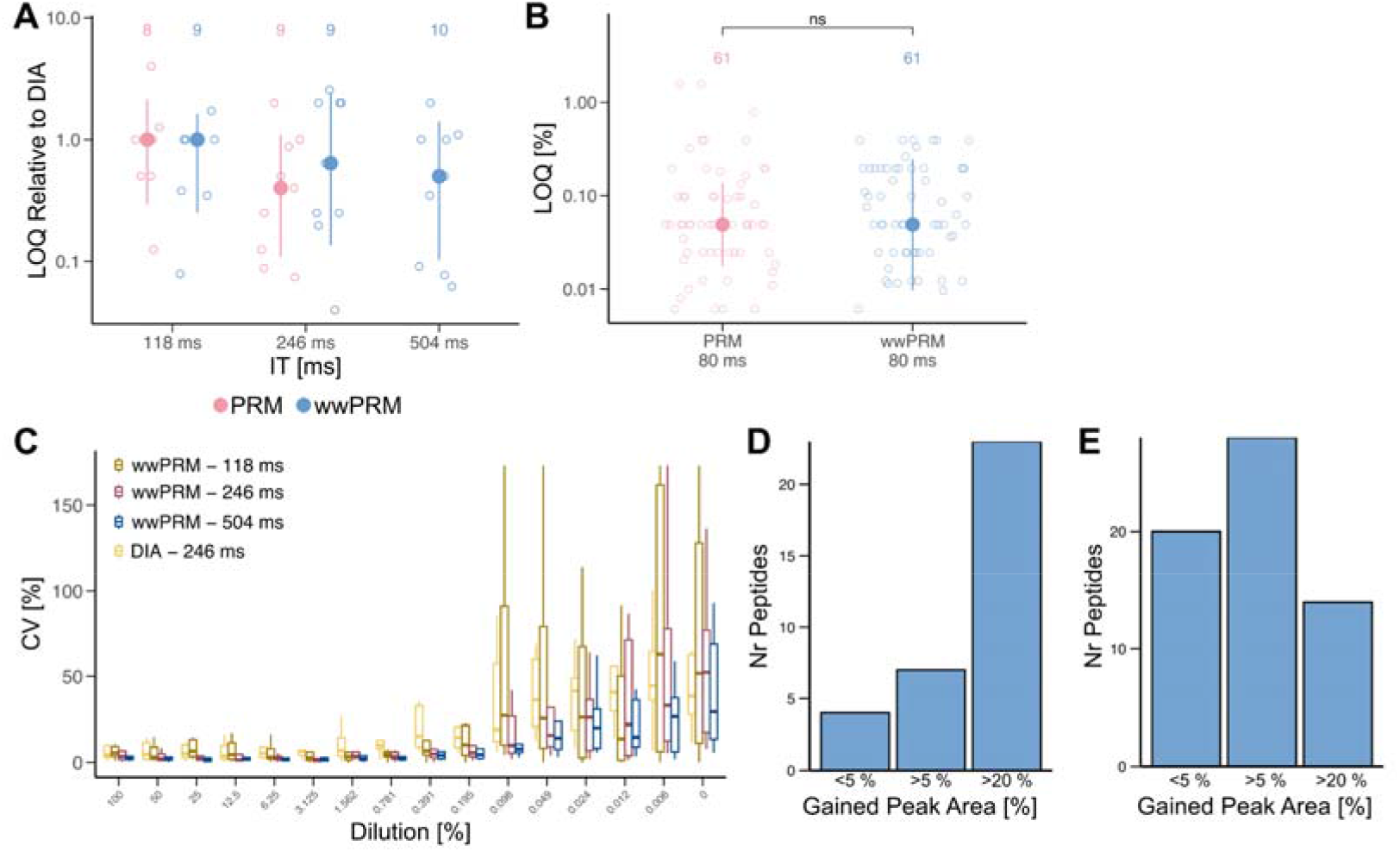
Increased MS2 isolation window width for targeted MS2 scans in low-input proteomics. (**A, C**) Quantitative comparison of PRM and wwPRM acquisitions on an Orbitrap Eclipse using a two-fold dilution curve of 10 synthetic peptides over 16 points in a 250 pg Hela digest, mean and standard deviation shown. Quantification based on ratio of light peptide to SIL standard. (**A**) LOQ of peptides acquired in PRM and wwPRM relative to their respective LOQ in DIA. (**B**) Quantitative comparison of PRM and wwPRM acquisitions on a Orbitrap Astral Zoom using a dilution curve of 61 synthetic peptides over 16 points in a 100 pg yeast and e. coli digest, Median and MAD shown. T-test corrected by Benjamini & Hochberg (**C**) Coefficient of variation (CV) of 10 peptides quantified in triplicates at each point of the dilution curve in DIA and wwPRM. Maximum injection times of 118 ms, 246 and 502 ms were used. (**D**) Percentage MS2 peak area gained with optimized NCE and FAIMS CV settings for 61 peptides compared to DIA settings. Optimized parameters used in data for Figure 1. (**E**) Percentage MS2 peak area gained with optimized NCE and FAIMS CV settings for 34 peptides compared to DIA settings. *** p < 0.001, ** p < 0.01, * p < 0.05, ns p > 0.05

**Supplementary Figure 3.**
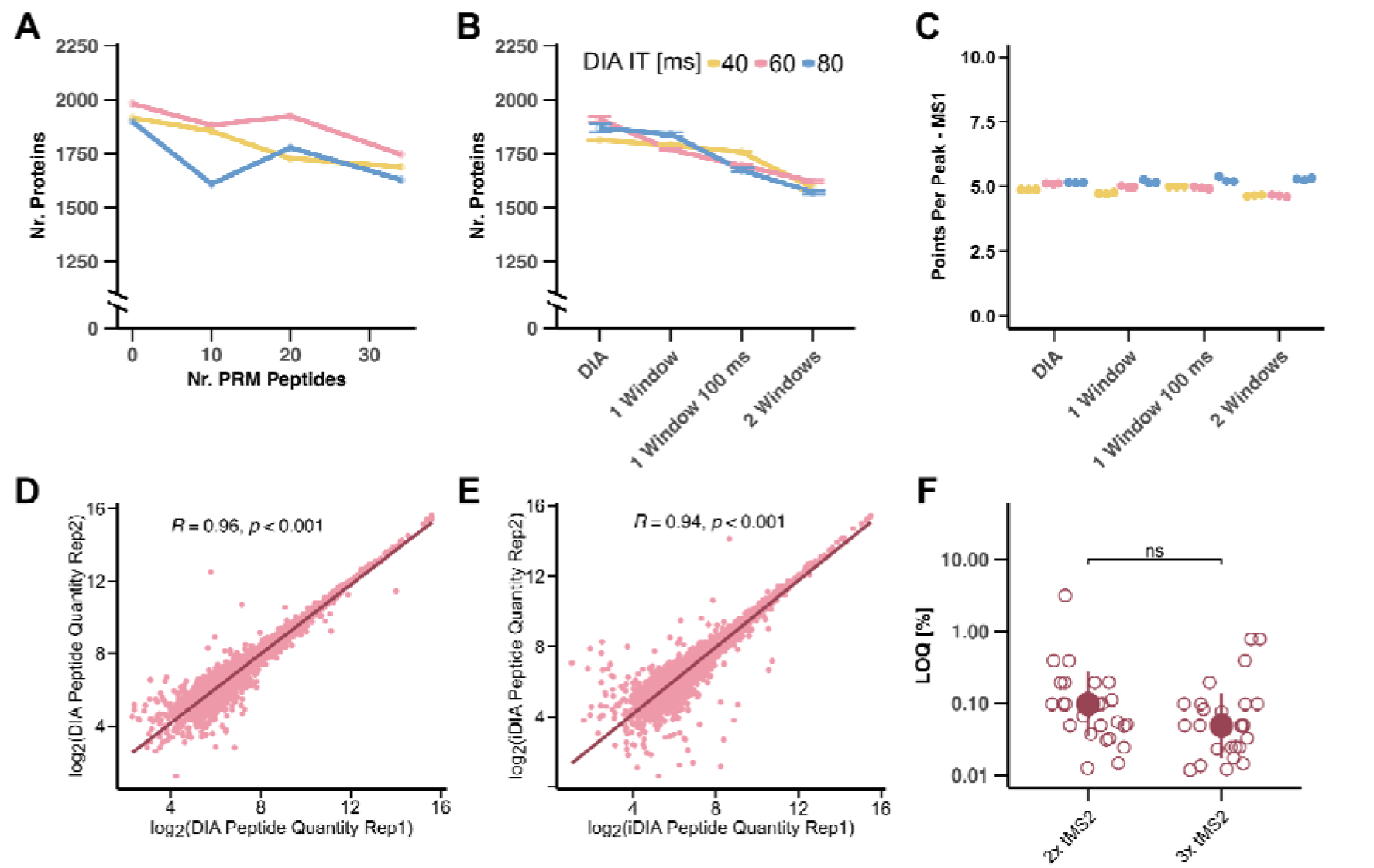
iDIA enables balance between global and targeted proteome coverage. 34 SIL peptides were spiked into 50 pg HeLa digest and acquired with iDIA methods based on three DIA methods displayed in Supplementary Table 1. (**A**) Number of proteins identified in triplicate acquisition of iDIA method quantifying 10, 20 or 34 peptides with wwPRM scans compared to DIA. (**B**) Number of proteins identified in triplicate acquisition of iDIA methods shown in Figure 2C. (**C**) MS1 points per peak in iDIA compared to DIA at three iDIA settings. Replacement of a DIA window with a wwPRM scan, replacement and extension of maxIIT of a DIA window with a wwPRM scan or replacement of two DIA windows by one wwPRM scan. (**D**) Pearson correlation or peptides between two representative DIA acquisitions at 60 ms maxIIT with 20 windows. (**E**) Pearson Correlation or peptides between two representative iDIA acquisitions at 60 ms maxIIT with 20 windows. (**F**) LOQ of 34 SIL peptides in wwPRM. *** p < 0.001, ** p < 0.01, * p < 0.05, ns p > 0.05

**Supplementary Figure 4.**
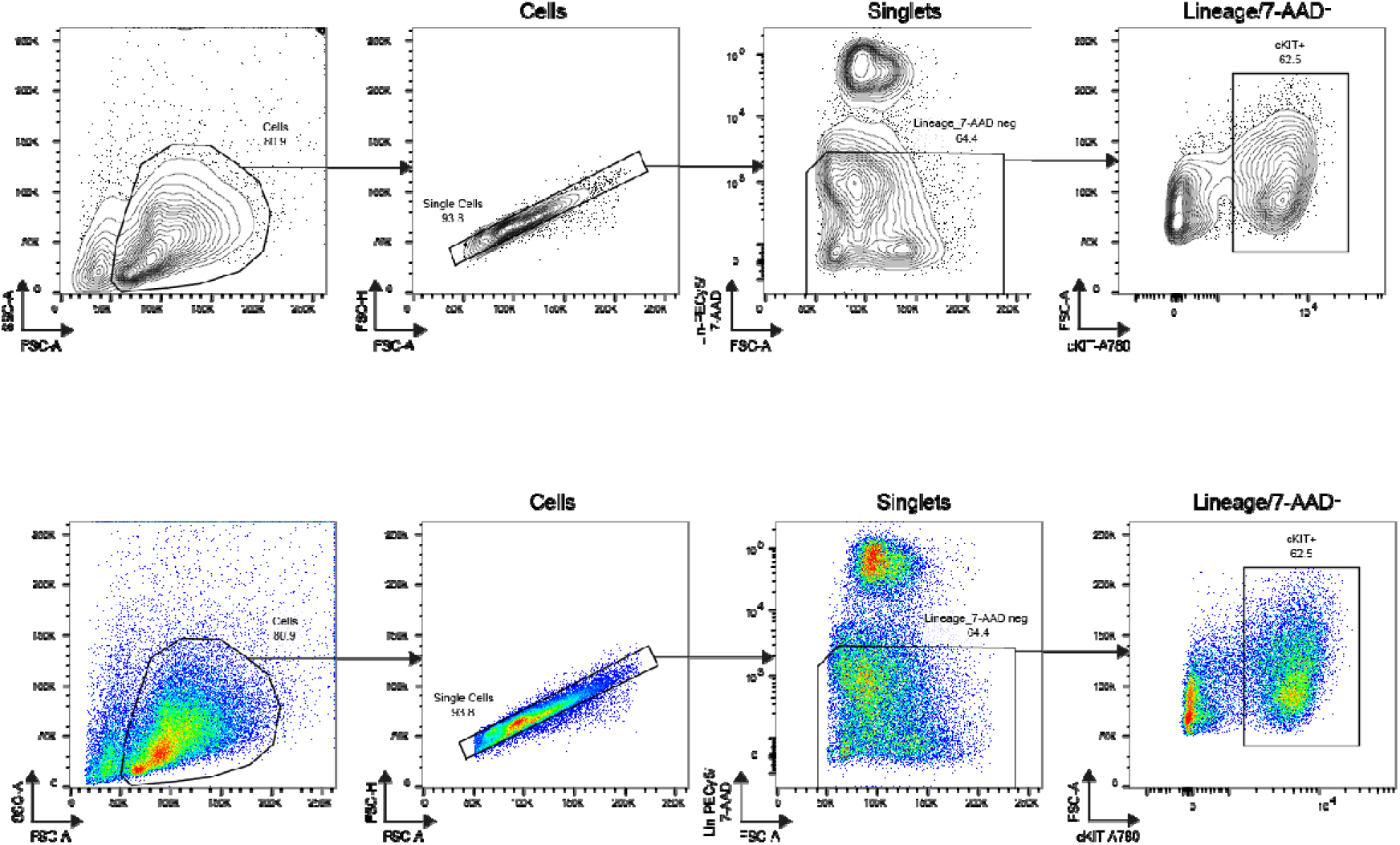
FACs Isolation schema of cKIT+Lin– mouse bone-marrow single cells.

**Supplementary Figure 5.**
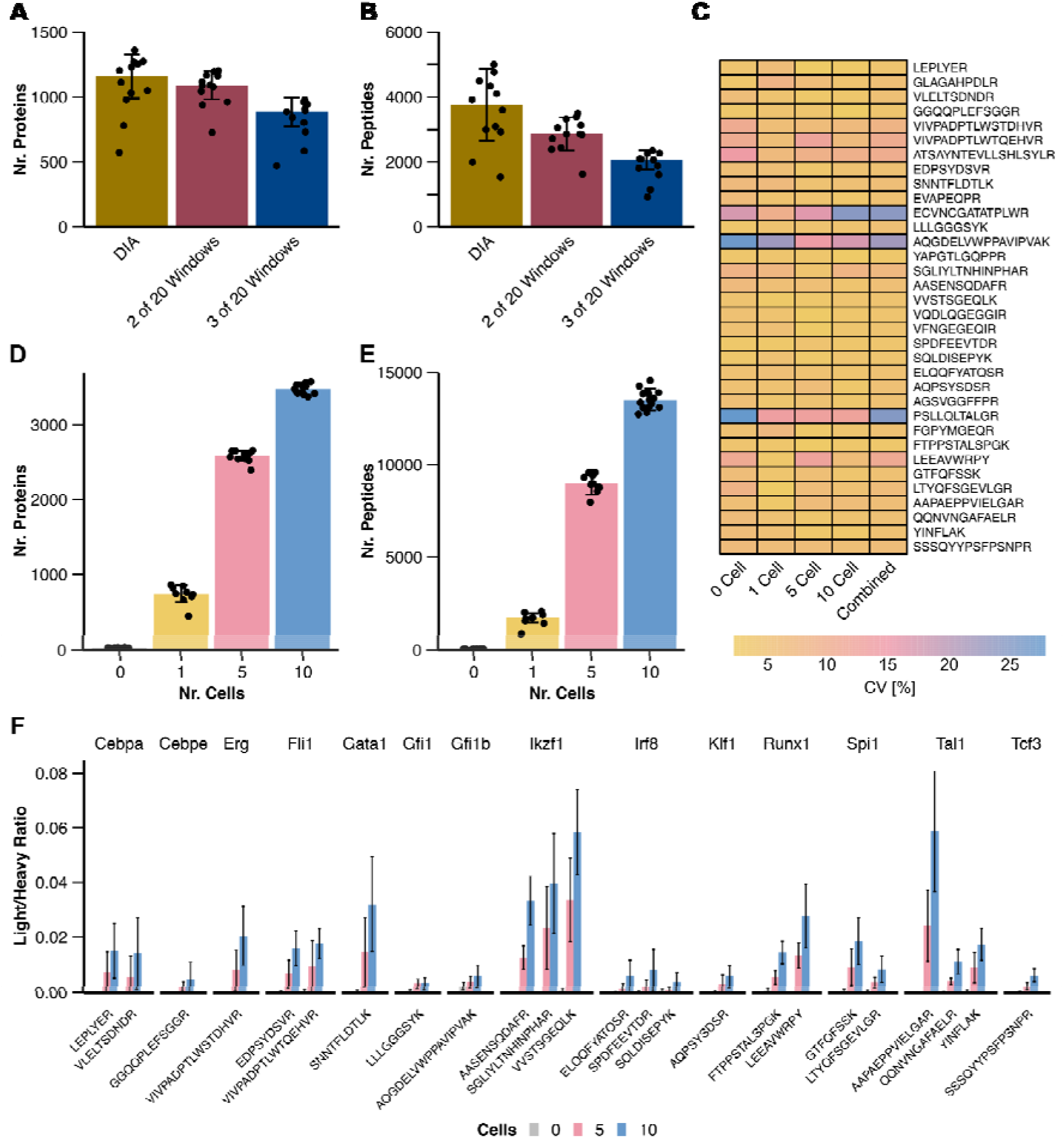
iDIA enables quantification of TFs in single and pooled 5 and 10 mouse HSCP cells. (**A, B**) Number of peptides and proteins identified from single cKIT+ cells with DIA and iDIA. Both methods operated DIA MS2 scans at 60 ms max IIT, 20 m/z DIA windows and 20 windows with a loop control of 0.6 seconds. iDIA replaced 2 or 3 DIA windows for each of 34 peptides per cycle with wwPRM scans. (**C-F**) 0, 1, 5 and 10 cKIT+ cells acquired in iDIA targeting 34 peptides covering 19 TFs. (**C**) CV of SIL raw peak area of the top 3 fragments for each peptide. (**D**) Proteins and (**E**) Peptides identified in 0, 1, 5 and 10 cKIT+ cells, Median and Mad displayed. (**F**) Ratio between top 3 fragments for each endogenous to SIL peptide for identified TF peptides, median and mad displayed.

**Supplementary Figure 6.**
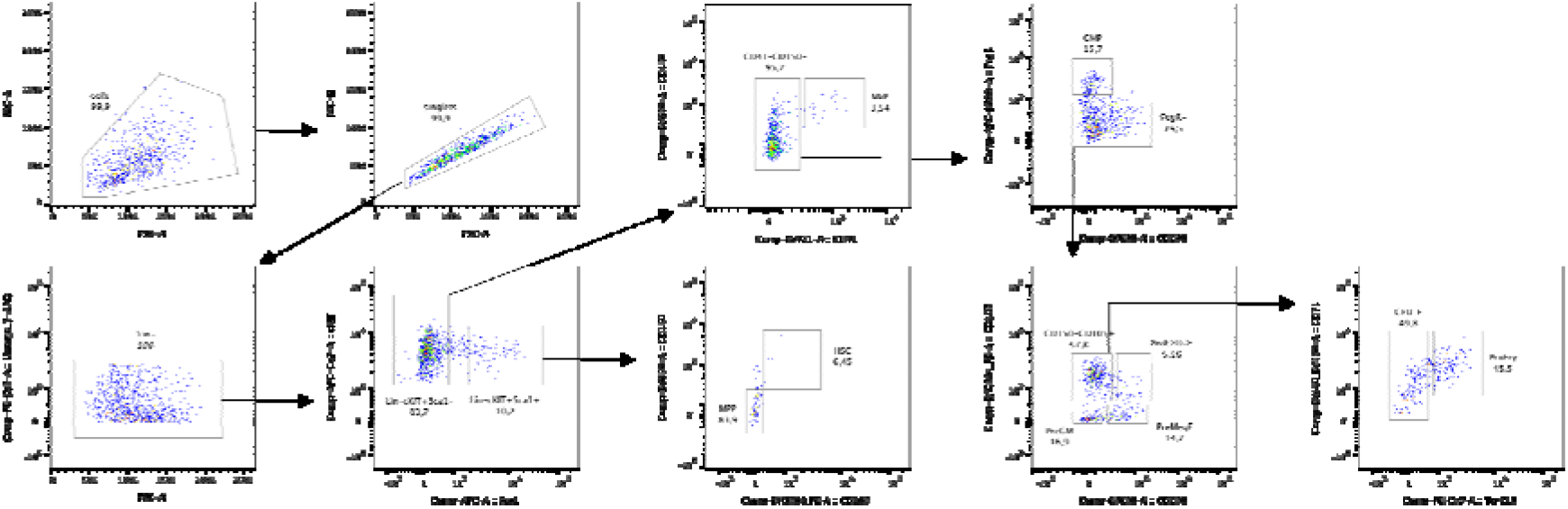
Cell-type annotation of index sorted cKit+Lin– bone-marrow cells. Sorted cells were retrospectively gated based on antibody-stained facs signals according to Pronk et. al. 2007 to visualize cell types.

**Supplementary Figure 7.**
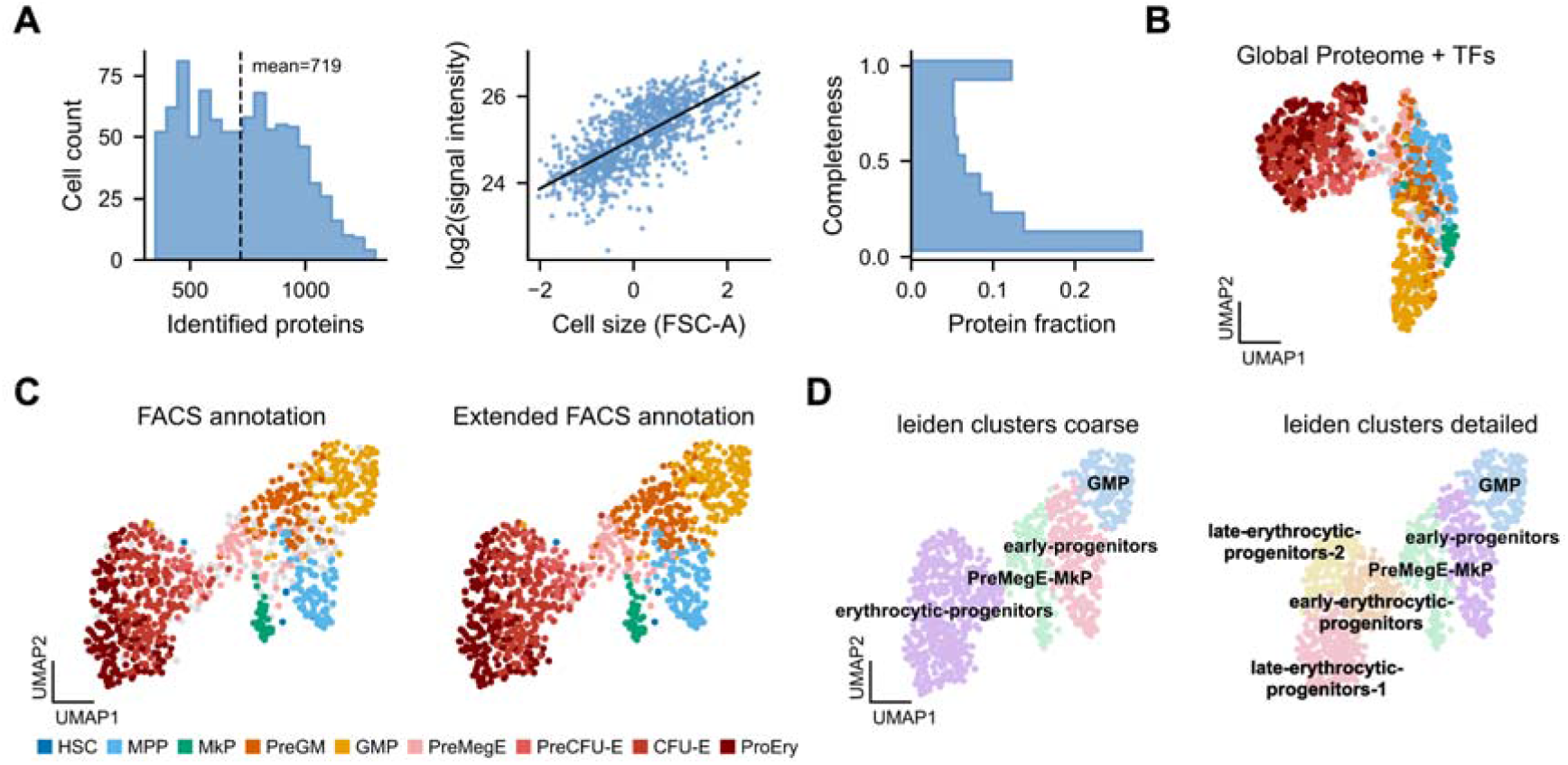
cKIT iDIA dataset overview. **(A)** Dataset summary metric **(B)** combined WNN embedding of Global proteome and TFs **(C)** UMAP of combined embedding highlighting the label transfer of FACS based cell-type annotation of unannotated cells using label-spreading. **(D)** Annotated leiden clusters at 0.4 and 0.6 resolution.

**Supplementary Figure 8.**
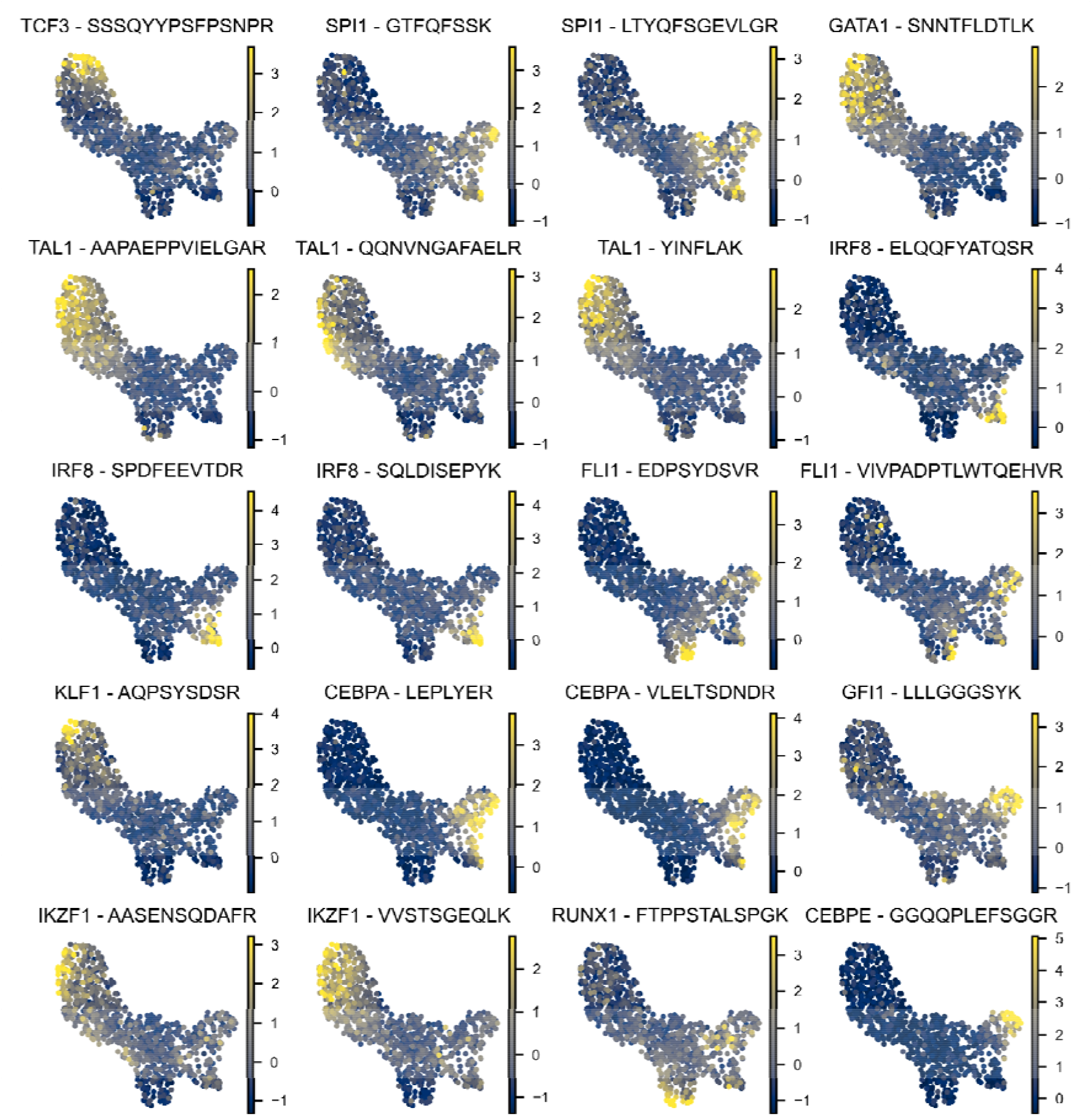
UMAP of TF expression overlaid with z-scored TF abundance.

**Supplementary Figure 9.**
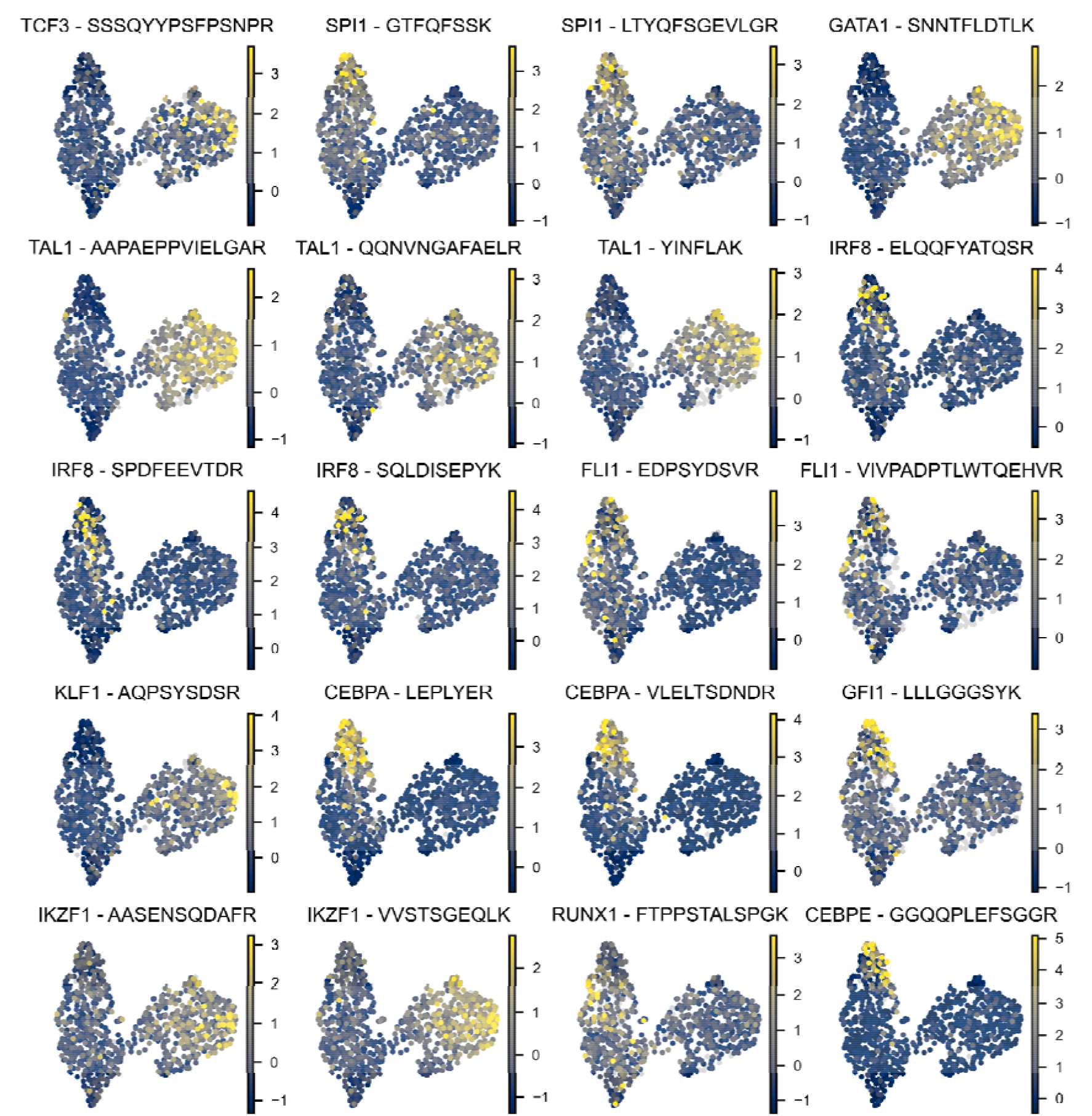
UMAP of global proteome expression overlaid with z-scored TF abundance.

**Supplementary Figure 10.**
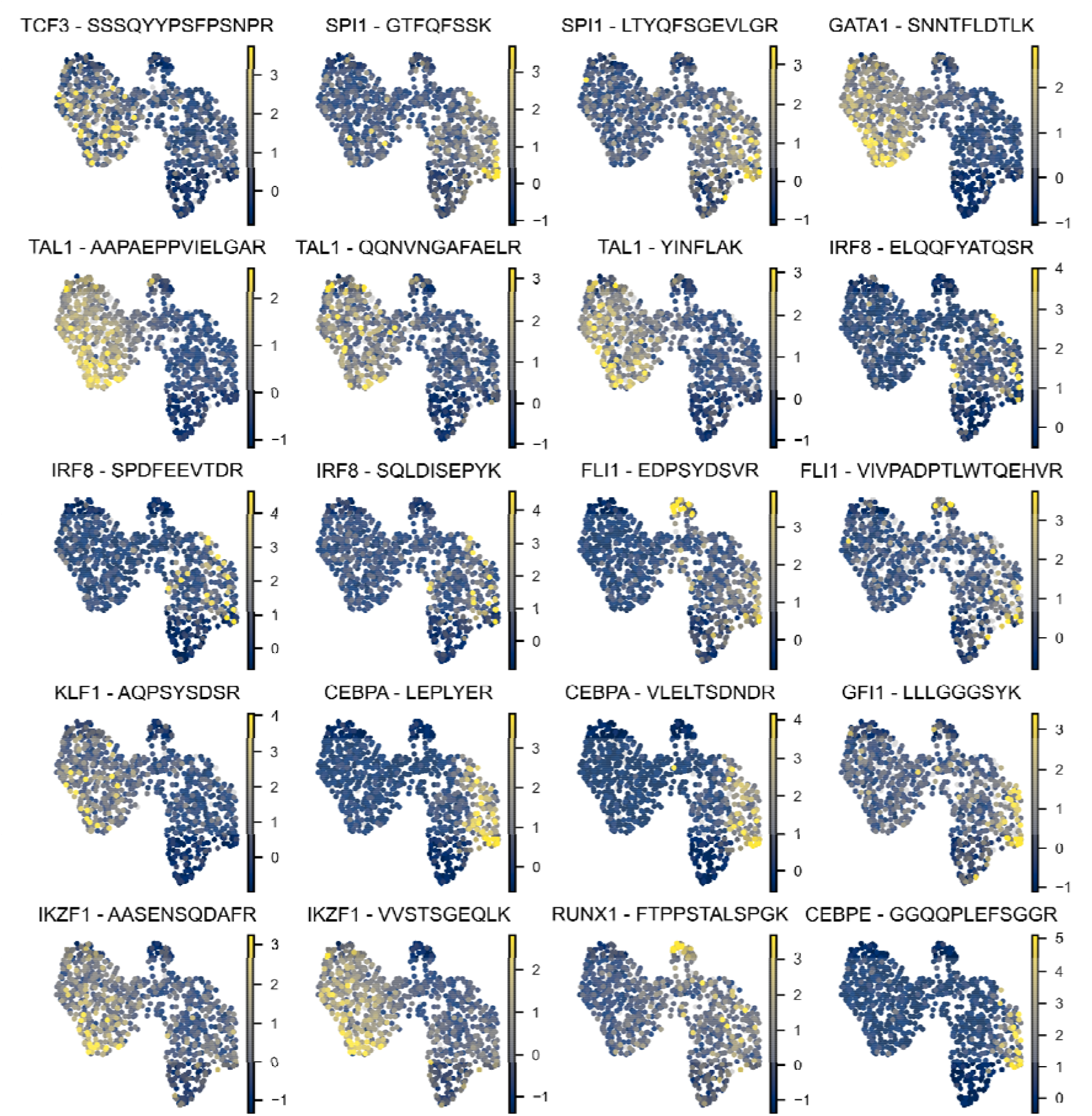
UMAP of FACS marker expression overlaid with z-scored TF abundance.

**Supplementary Figure 11.**
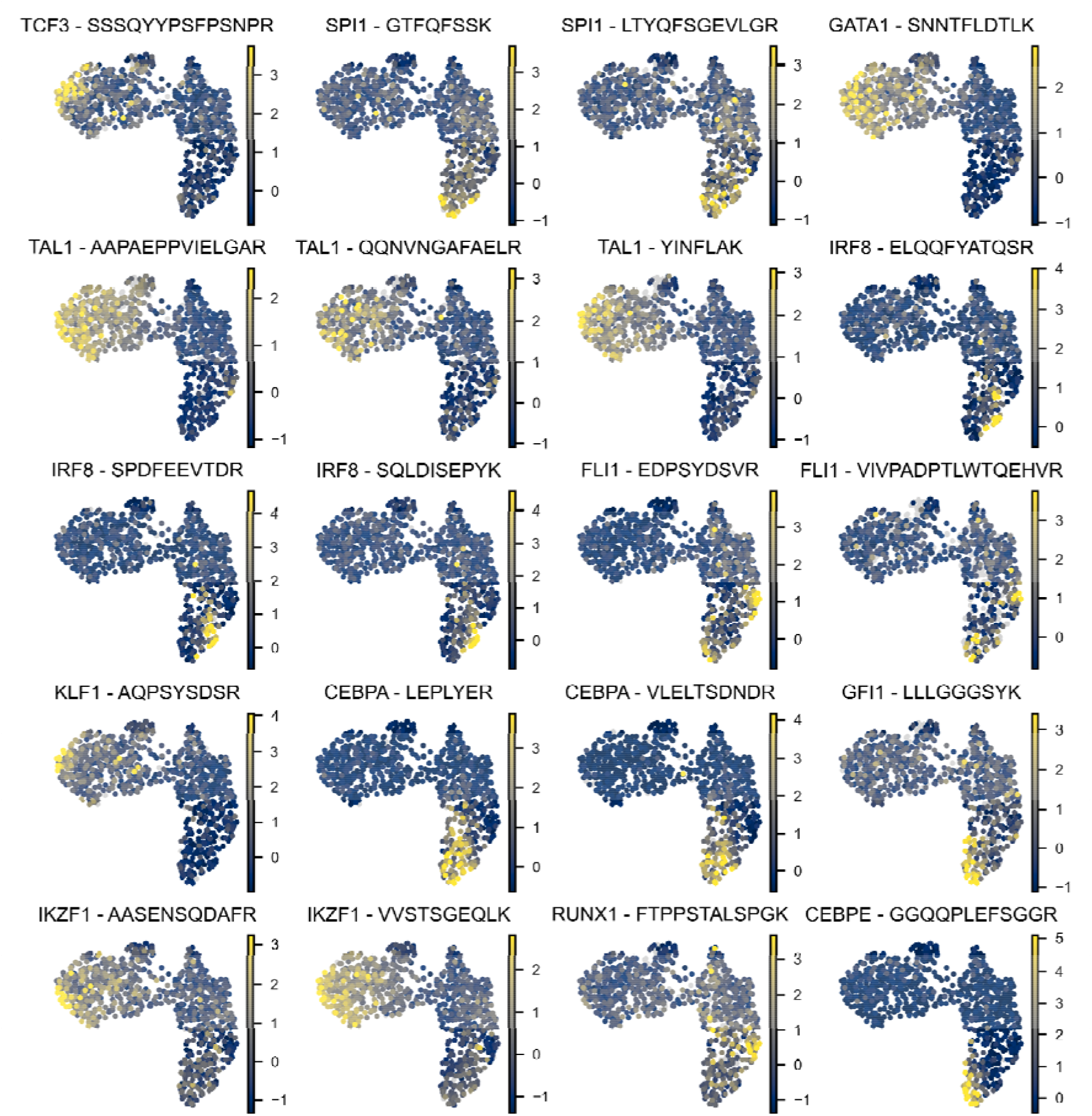
UMAP of multimodal embedding of global proteome and TF overlaid with z-scored TF abundance.

**Supplementary Figure 12.**
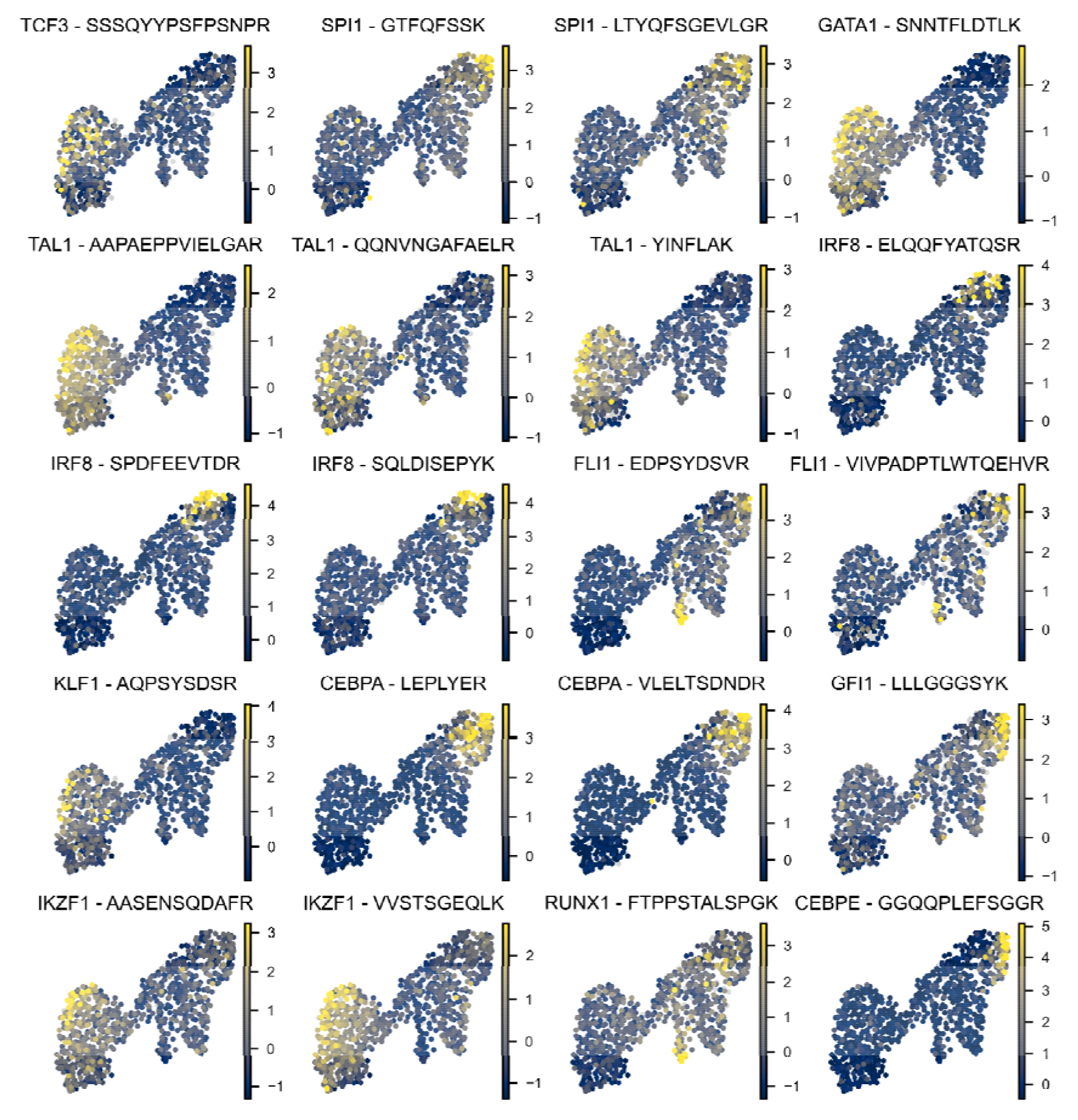
UMAP of multimodal embedding of global proteome, TF and FACS markers overlaid with z-scored TF expression.

**Supplementary Figure 13.**
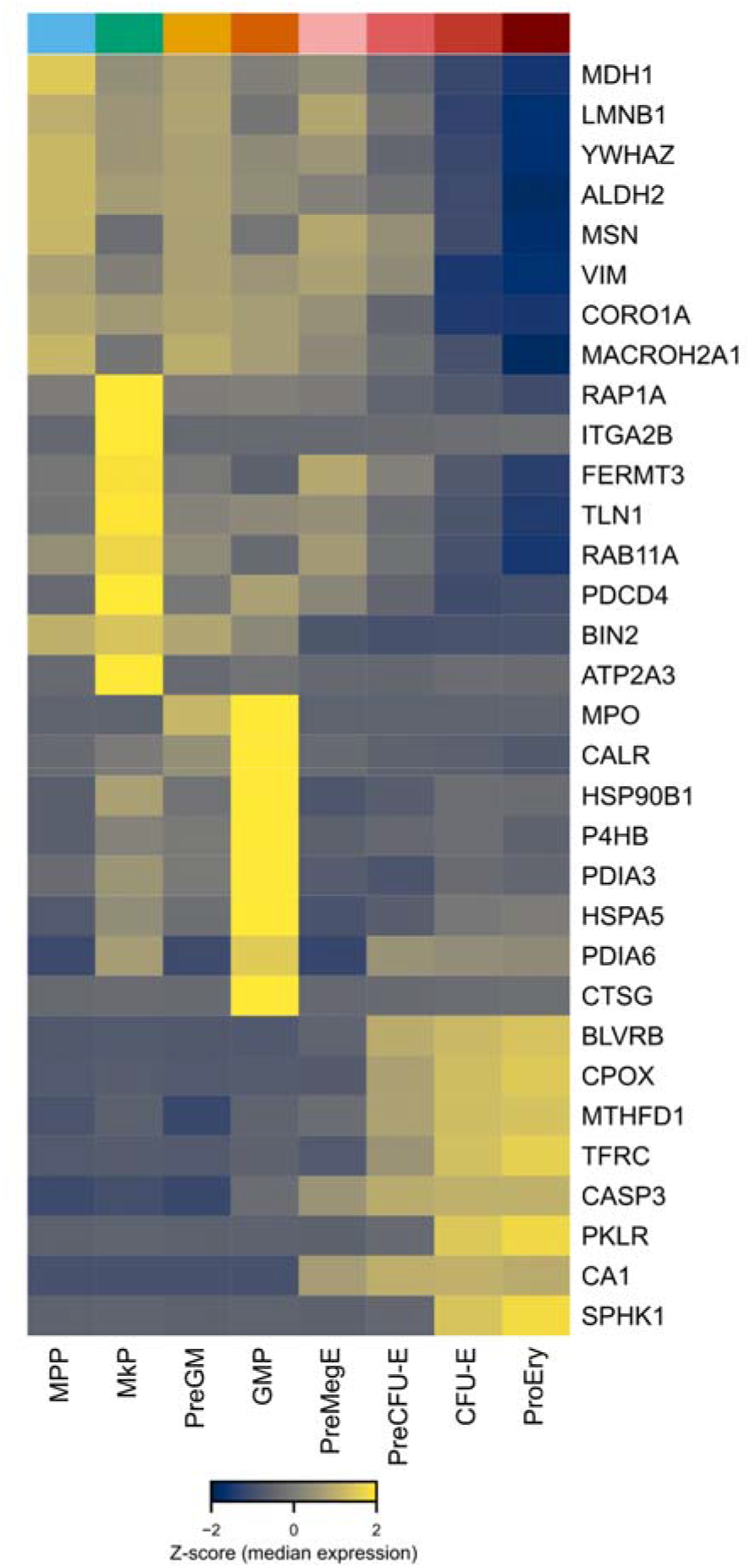
Cell-type specific ranked gene expression for GMP, Ery and HSC/MPP.

**Supplementary Figure 14.**
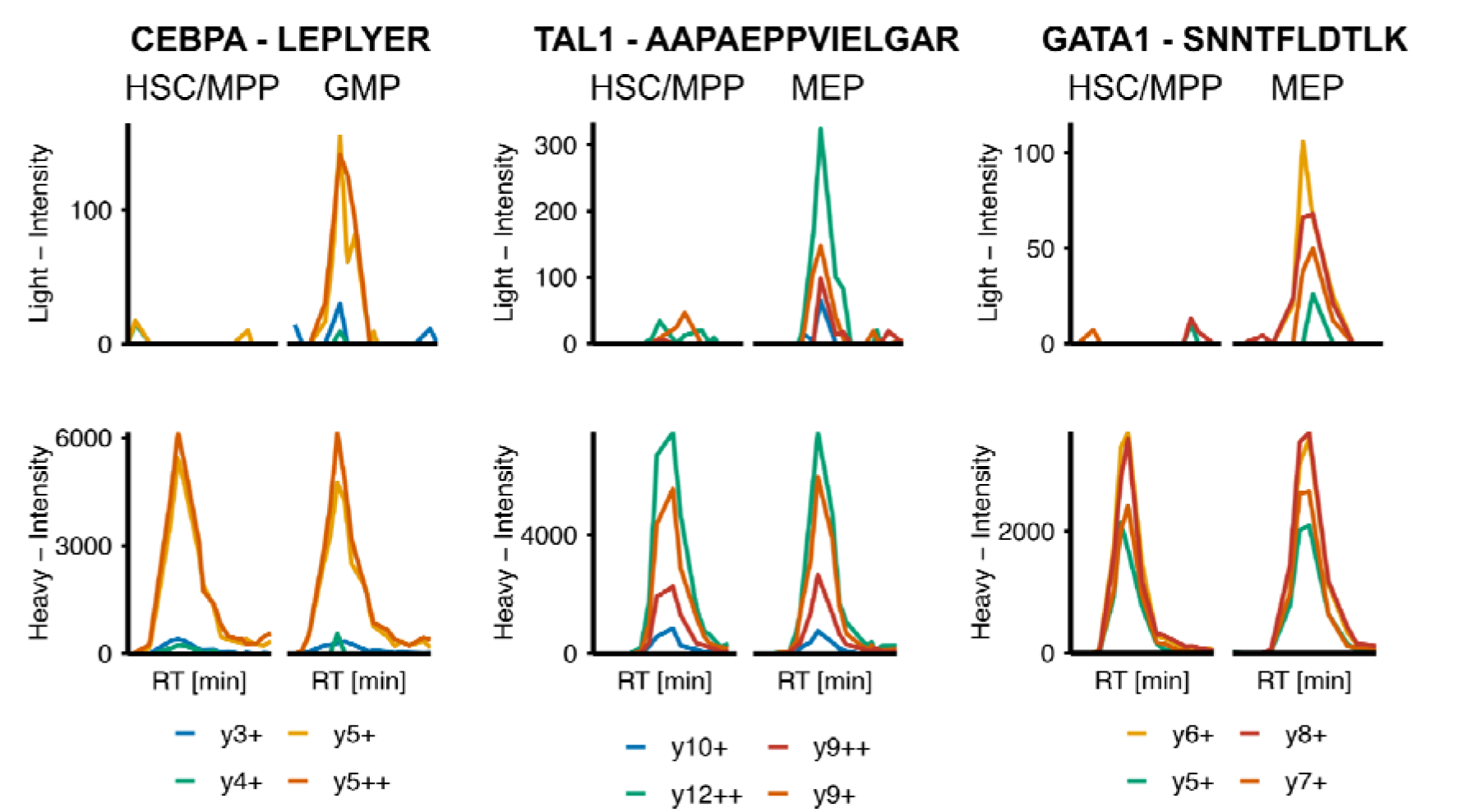
Detailed XICs corresponding to Figure 3E. Example HSC/MPP cells were selected based on the top 10 lowest cells in pseudotime with the cell closest to mean expression for the illustrated TF. Erythroid-megakaryocyte progenitors and GMP example cells were manually selected of the 20 cells with the highest quantity of the respective cell-type (GMP or ProEry, CFU-E).

**Supplementary Figure 15.**
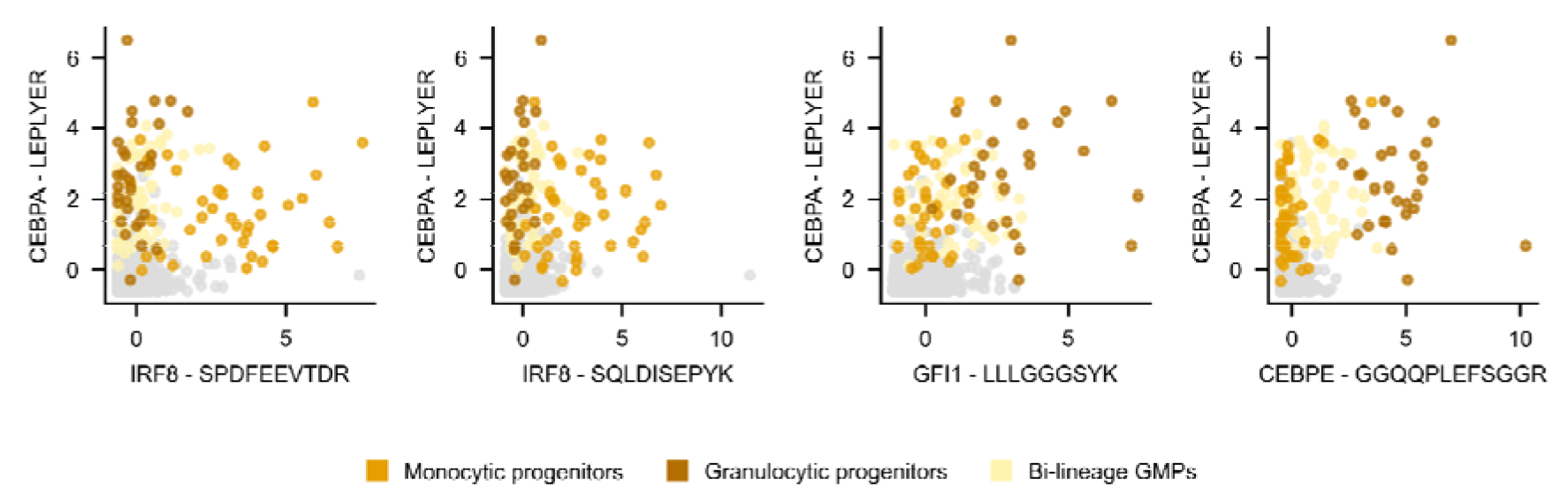
Correlation of CEBPA expression with IRF8, GFI1 and CEBPE in GMP subpopulations. Cells not part of the leiden GMP cluster are displayed in grey. Z-scored TF abundance displayed for each TF peptide.

## Methods

### Cultivation of HEK 293-F cells

Hek 293-F cells were cultured at 37 °C in Gibco™ FreeStyle™ 293 Expression Medium containing Penicillin and Streptavidin. Cells were washed three times in PBS prior to sorting.

### Isolation of cKIT-Cells

One 13-week-old female C57BL/6 mouse was sacrificed by cervical dislocation and leg bones (femur, tibia and iliac crest) and spine were isolated. The bones were crushed three times with 10 mL PBS+3% FBS and the suspension was filtered through a 70μm cell strainer. The filtered suspension was then centrifuged at 300g for 10 min at 4°C. To enrich for cKit+ (CD117+) cells, the resulting pellet was resuspended in 194µl PBS+3% FBS and 6µl CD117 MicroBeads (Miltenyi Biotec, 130-091-224) and incubated on ice for 30 min. Magnetic separation was carried out using an LS column (Miltenyi Biotec, 130-042-401), according to manufacturer’s protocol.

### FACS sorting

Following cKit enrichment, the BM cells were stained with antibodies on ice for 30 min (see **Supplementary table 3**). Following staining, the cells were washed three times with PBS to remove any residual serum by centrifugation at 300g for 10 min at 4 °C. The pellet was resuspended in PBS containing 1:1000 (v/v) 7-AAD viability dye (Invitrogen, A1310). Using a BD FACSymphony S6 cell sorter with a 100-micron nozzle set to single-cell sort mode, one cell was sorted into each well of an Eppendorf twin.tec 384 LoBind plate containing 1µl of 20% TFE, 80mM TEAB lysis buffer. Immediately after the sort, plates were briefly spun down, snap-frozen on dry ice, and stored at –70°C.

### Sample Preparation

Single cells were sorted into a 384 well plate containing 1 µL of lysis buffer (80 mM Triethylammonium bicarbonate (TEAB) pH 8.5, 20% 2,2,2-Trifluoroethanol (TFE)). The plate was frozen to –80 °C. Prior to sample acquisition the plates were heated to 95 °C followed by a second freezing cycle to –80 °C. 1 µL containing 2 ng Trypsin Platinum (Promega) was added to each well and the plate was incubated at 37 °C overnight. The digestion was stopped by the addition of 1 % trifluoroacetic acid (TFA). If SIL peptides were used in the experiment they were diluted to their final concentration in 1 % TFA and added to the single cells during the quenching of the digestion.

Bulk samples were prepared from Pierce™ Yeast Digest Standard, Pierce™ HeLa Protein Digest Standard or MassPREP E.coli digestion standard (Waters™). Standards were reconstituted in 0.1 % Formic Acid (FA) and diluted to the respective concentration in 0.1 % FA.

### Peptide Standards

Peptide standards were ordered from JPT Peptide Technologies GmbH. Upon receival peptides were resuspended in 0.1 % FA and stored at –80 °C. Peptides concentrations were adjusted to the respective sample matrix in an iterative process.

### LC-MS analysis

Samples were analyzed using the Vanquish Neo UHPLC system (Thermo Scientific) in combination with the Thermo Scientific Orbitrap Astral and a Thermo Scientific Orbitrap Astral Zoom MS as well as the Ultimate 3000 nano-LC (Thermo Fisher Scientific) combined with the Thermo Scientific Orbitrap Eclipse. All instruments were equipped with FAIMS Pro interface (Thermo Scientific) or FAIMS Pro Duo interface (Thermo Scientific) in combination with the Easy-Spray Source.

Peptide separation was performed on the 50 cm µPAC Neo analytical column and the 50 cm µPAC Neo Plus analytical column combined with the µPAC Standard Trap Columns and the EASY-Spray™ Emitters (Thermo Scientific).

On the Orbitrap Astral MS1 spectra were acquired with resolution of 240,000, a maximum injection time of 100 ms and the automated gate control (AGC) was set to 500 %. During DIA acquisitions MS2 scans were performed at an AGC of 500 % over the mass range of 400 m/z to 800 m/z at a HCD collision energy of 25. Maximum injection time (max IIT), DIA-window width and Loop Control were set to complete one full DIA cycle in approximately 1.2 seconds with a Loop Control 0.6 seconds at max IIT. Injections times of 40 ms, 40 ms, 60 ms and 80 ms were combined with a respective DIA-window width of 13.4 m/z, 13.7 m/z, 20 m/z, 26.6 m/z and a loop control (N) of 16, 15, 10, 8. FAIMS was operated at a compensation voltage (CV) of –48 V.

PRM MS2 scans on the Orbitrap Astral were performed at 1.7 m/z isolation width at 40 ms and 80 ms max IIT. wwPRM scans were performed at an isolation width of delta(Heavy Isotope Precursor Mass – Light Isotope Precursor Mass)/ precursor charge state + 1.7 m/z with an injection time of 80 ms. AGC was set to 500 %. Both FAIMS CV and HCE collision energy were operated at custom settings for the respective peptides. FAIMS CV was optimized between –30 V and –55 V and HCD was optimized between 20 and 33.

iDIA acquisitions were performed with MS1 scans as stated above. MS2 scans were operated at an AGC of 500 %. DIA windows were acquired with an HCD collision energy of 25 and a FAIMS CV of –48 V over the mass range of 400 m/z to 800 m/z. Windows and injections time were as described for the DIA methods above. wwPRM scans were acquired with custom HCD and FAIMS settings at max IIT’s of 40 ms, 60 ms, 80 ms, 100 ms, 120 ms or 160 ms. The loop control was set to 0.6 seconds or N of 16, 15, 10 or 8. wwPRM scans were performed over 0.5 min for each target peptide. Acquisitions performed on the Orbitrap Astral Zoom were done with the Low Input Application Mode enabled.

On the Orbitrap Eclipse MS1 spectra were acquired at a FAIMS CV of –45 V at a Orbitrap resolution of 120,000 and a max IIT of 246 ms. AGC was kept at 300 %. DIA MS2 scans were acquired at a resolution of 120,000 or 240,000 with a max IIT of 246 or 502 ms. DIA windows were acquired at 68 m/z width over the mass range of 400-800 m/z. AGC was kept at 1000 % with a collision energy of 26 and a FAIMS CV of –45 V. PRM acquisitions were performed at an isolation window width of 1.7 m/z with custom collision energy and FAIMS CV settings. AGC was kept at 300 %. PRM scans were acquired at resolutions of 60,000, 120,000 and 240,000 with the respective max IIT of 118 ms, 246 ms and 502 ms. wwPRM MS2 scans were operated at an AGC of 300 % with custom FAIMS CV and collision energy settings. Isolation windows were set to delta(Heavy Isotope Precursor Mass – Light Isotope Precursor Mass)/ precursor charge state + 1.7 m/z. iDIA acquisitions were performed at a MS1 resolution of 120,000 with a maximum IIT of 246 ms. The AGC was kept at 300 % and FAIMS CV was set to – 45 V. MS2 scans were operated in a combination of DIA and wwPRM scans. DIA scans were performed over the mass range of 400-800 m/z with 68 m/z wide MS2 windows. Collision energy and AGC were set to –45 V and 300 % respectively. wwPRM scans were operated as described above with a custom FAIMS CV and collision energy for each peptide. wwPRM scans were operated over the time of 0.7 minutes for each target peptide with a loop control of N=3.

For the 50 cm µPac Neo Plus analytical column the gradient was built at 600 nL/min or 750 nL/min starting at 1 % Solvent B with an increase to 10 % solvent B over the first 0.05 minutes. During the following 2.1 minutes Solvent B was increased to 22.5 %. The next 1.25 min Solvent B was increased to 37.5 % followed by an increase to 40% within 0.1 min. Solvent B was further increased to 90 % for the remainder of the gradient. At minute 3.5 of the gradient the flow was dropped to 200 nL/min. Samples were acquired over 12.5 min, 12.6 min and 13.5 min. The column was kept at 50 °C.

The 50 cm µPac Neo analytical column was operated at 50 °C. The column was loaded at 750 nL/min at 100 % Solvent A. After 0.2 min Solvent B was increased to 8 % followed by a linear increase to 24 % B until minute 3 at which point the flowrate was dropped to 200 nL/min. The linear gradient was raised to 48 % B at minute 8 and increased to 99 % B over the following 0.4 minutes. 99 % B was kept for 6.55 minutes followed by a drop to 1 % B until minute 17.85.

### PRM development

Synthetic peptides were acquired in DIA, and the identified precursors were subjected to a scheduled PRM scan. PRM scans were manually revised followed by PRM acquisitions at FAIMS CVs between –30 V and –55 V and HCDs between 20 and 33. Notably the MS1 scans were kept at stable FAIMS CV –45 V (Eclipse) and –48 V (Astral) and stable HCD. Only FAIMS CV and HCD of the MS2 scans were altered. The total MS2 peak area of each peptide was normalized to the MS1 peak area to identify the optimal FAIMS and HCD settings for each peptide. A peptide library was built from these optimized settings containing the top 15 b-and y ions >b2 and >y2. To address the quantitative performance of these PRM assays, standard curves were established of the synthetic peptides in a stable background matrix of 250 pg, 100 pg or 50 pg Hela digest or combined Yeast and E coli digest.

## Data Analysis

### LOQ determination

Standard curves were established by performing a serial dilution of the synthetic peptides in a stable sample matrix as described above. Peptide standards were titrated into the respective matrix to start each standard curve at the highest protein expressed in the background proteome. Peptide standards were diluted over 14 to 16 dilution points with a sample containing the matrix as the last point of the standard curve to model the background signal of each peptide.

Results were imported into Skyline-daily ^27^ (version 25.0.9.97) and peptide peak integration was revised. All b, y and precursor ions that were part of the library were exported at fragment level peak area. The fragment level elution profile was compared to the library, and the most intense top 3 y-fragment ions were selected for quantification. B-ions were only used for identification of the peptides and peak area integration. In case of wwPRM acquisitions the b-ions would be shared between heavy and light peptide and were therefore not considered for quantification. The top three y-ion peak areas of each heavy as well as light were summed. In case both heavy and light were used for quantification the ratio of the summed peak-area between heavy and light was taken. In case only light or only heavy peptides were quantified, the summed peak area was normalized to the total ion current (TIC) of the respective run. The CV between the replicates of each point of the standard curve were calculated and only replicates with a CV below 20 % were considered to fit the standard curve. The LOD of each peptide was identified by integrating the background matrix without peptide spike-in at the same retention time window as the background matrix spiked with synthetic peptides. The LOD was defined as mean(sum peak area background matrix) + 2*Standard Deviation (sum peak area background matrix). To allow for a noise reduction during the fitting of the standard curve, replicates with a deviation from the expected linear ratio between dilution points were excluded if they had a delta variation larger then 25 %. The remaining dilution points were used to fit a linear model. Following assumptions were made when calculating the LOQ: If the fitted linear model contains the last dilution curve point above the LOD and no dilution curve point below the LOD has a replicate with a higher peak area than the LOD, the LOD is considered the LOQ. If this assumption was not met, the last point of the dilution curve that was used to fit the linear model is considered the LOQ.

### iDIA setup

To compile an iDIA MS method, the preferred DIA window setting was exported from the Method Editor. Additionally, the established PRM target list containing the heavy or light version peptides of interest, their mass, retention time window, optimal FAIMS CV and optimal HCD was exported from the method editor. The sample of interest was acquired with the preferred DIA method and searched in Spectronaut 19.9. The precursor level search results were exported containing the m/z of each precursor and its peak area start and end. These three csv files were passed to the iDIA generator together with the information of the base peak width of the used chromatography, cycle time of the DIA method, DIA NCE and FAIMS CV as well as DIA-MS2 IIT and preferred PRM IIT. PRM methods were converted to wwPRM method if specified and the replacement of more than one DIA window with PRM scans could be specified. The iDIA method generation considered the number of precursors that were identified in each combination of DIA windows that were to be replaced by PRM scans and selected the combination of DIA windows that contained the least precursors while maintaining an equal time gap between the PRM acquisitions. PRM scans were converted into wwPRM scans which include both heavy and light peptide with a 0.85 m/z spacing before and after the light and heavy precursor. DIA windows were replaced with wwPRM scans at the specified retention time window in the PRM method. A csv file containing a combined iDIA method that contains both scheduled DIA and wwPRM scans over the full chromatographic gradient was imported into the method editor.

### DIA Search Settings

DIA and iDIA raw files were searched in both DIANN 2.1 and Spectronaut 19.9. Depending on the sample matrix, Carbamidomethyl (C) was set as a fixed modification. DirectDIA was used in Spectronaut with the respective FASTA file. Quantification was performed on the MS1 level with the FDR set to 0.01. If multiple conditions were searched together, each condition was set, and Method Evaluation was selected. In DIANN 2.1 a library was built based on the FASTA file using DIANN. Trypsin/P with 1 missed cleavage was selected and N-term Methionine excision enabled. Peptide length was set to 6-30 AA and precursor range was specified as in the DIA methods ranging from 400 m/z to 800 m/z. The FDR was set to 1 %.

### Single Cell Data Processing

Raw files were searched with DIANN 2.1 using MBR against the mouse swissprot proteome (version: 30.04.2025) as well as a mouse specific contamination fasta file ^28^. Single cell raw files and negative control (sample without sorted cell) raw files were imported into Skyline. Only MS2 spectra with a m/z window covering less than 7 m/z were imported. Peptides were further specified to be only extracted from their unique wwPRM scans. Peak area integration was performed by Skyline on the SIL peptides. A fragmentation level report was exported. Results were further processed in R and python.

Based on the DIA-global proteome cells were excluded from the analysis if they had below 350 identified proteins in the global proteome and a low correlation of the total MS signal to FSC-A. Missing proteins were imputed with half of the minimum protein expression if present in >30 cells. Targeted proteomics TF data was further filtered. TF peptides were excluded in a first step based on the SIL peptide data quality. Peptides with a SIL peptides with a dotp < 0.75 and/or a total peak area (5 fragments) of less than 2000 and/or a total peak area < 1/4 of mean total area across all samples and/or a weight of less than 0.5 based on its contribution to a MM-type linear regression estimators fitted to the total area over time were excluded. Furthermore, samples with <50 % of the SIL peptides passing the above criteria were excluded. Next the negative control was used to identify high background signal and fragments with an area above 100 in over 50 % of the negative controls were excluded for quantification. Next an endogenous peptide was considered identified and quantified within the total dataset if over 1 % of the cells contained the peptide with a dotp above 0.75 (based on 5 fragments), less than 50 % of the cells contained only 1 fragment ion for the endogenous peptide (majority of cells has no expression of the TF), if >40 percent of the cells contain fragment peak areas above the negative control and the endogenous and SIL peptide displayed co-elution. Peptides were quantified by ratio to SIL peptide of the top two SIL fragment ions. Ratios based on an endogenous peptide peak area of 0 were modeled with a non-negative value below the smallest ratio above 0 for data visualization.

Cell-types were annotated based on antibody based facs gates and cells without annotation were further annotated by label spreading. A weighted nearest neighbor (WNN) ^17^ analysis was used to integrate global proteome, TFs and Facs cell surface markers into a joined embedding. Missingness in TFs only resulted from technical missingness, and the median 3.5 % missing data were imputed by knn-imputation. Imputed TFs were only used in Figure 3D and 5A, otherwise non-imputed data was used. For XIC visualization, scans with identical exported retention-time values due to Skyline rounding were separated by a 0.005 min offset to preserve plotting continuity.

## References

1. Furtwängler, B. et al. Mapping early human blood cell differentiation using single-cell proteomics and transcriptomics. Science 0, eadr8785 (2025).

2. Bubis, J. A. et al. Challenging the Astral mass analyzer to quantify up to 5,300 proteins per single cell at unseen accuracy to uncover cellular heterogeneity. Nat. Methods 22, 510–519 (2025).

3. Ye, Z. et al. Enhanced sensitivity and scalability with a Chip-Tip workflow enables deep single-cell proteomics. Nat. Methods 22, 499–509 (2025).

4. Petrosius, V. et al. Quantitative Label-Free Single-Cell Proteomics on the Orbitrap Astral MS. Mol. Cell. Proteomics 100982 (2025) doi:10.1016/j.mcpro.2025.100982.

5. Gillespie, M. A. et al. Absolute Quantification of Transcription Factors Reveals Principles of Gene Regulation in Erythropoiesis. Mol. Cell 78, 960–974.e11 (2020).

6. Peterson, A. C., Russell, J. D., Bailey, D. J., Westphall, M. S. & Coon, J. J. Parallel Reaction Monitoring for High Resolution and High Mass Accuracy Quantitative, Targeted Proteomics *. Mol. Cell. Proteomics 11, 1475–1488 (2012).

7. Eshghi, A. et al. Sample Preparation Methods for Targeted Single-Cell Proteomics. J. Proteome Res. https://doi.org/10.1021/acs.jproteome.2c00429 (2023) doi:10.1021/acs.jproteome.2c00429.

8. Martínez-Val, A. et al. Hybrid-DIA: intelligent data acquisition integrates targeted and discovery proteomics to analyze phospho-signaling in single spheroids. Nat. Commun. 14, 3599 (2023).

9. Xie, X. et al. Multicolumn Nanoflow Liquid Chromatography with Accelerated Offline Gradient Generation for Robust and Sensitive Single-Cell Proteome Profiling. Anal. Chem. 96, 10534–10542 (2024).

10. Nam, D. et al. Wideband PRM: Highly Accurate and Sensitive Method for High-Throughput Targeted Proteomics. Anal. Chem. 96, 10219–10227 (2024).

11. Demichev, V., Messner, C. B., Vernardis, S. I., Lilley, K. S. & Ralser, M. DIA-NN: neural networks and interference correction enable deep proteome coverage in high throughput. Nat. Methods 17, 41–44 (2020).

12. Picotti, P. & Aebersold, R. Selected reaction monitoring–based proteomics: workflows, potential, pitfalls and future directions. Nat. Methods 9, 555–566 (2012).

13. Hatton, I. A., et al. The human cell count and size distribution. Proc. Natl. Acad. Sci. 120, e2303077120 (2023).

14. Üresin, N. et al. Unraveling the proteome landscape of mouse hematopoietic stem and progenitor compartment with high sensitivity low-input proteomics. 2024.05.03.592307 Preprint at 10.1101/2024.05.03.592307 (2024).

15. Paul, F. et al. Transcriptional Heterogeneity and Lineage Commitment in Myeloid Progenitors. Cell 163, 1663–1677 (2015).

16. Pronk, C. J. H. et al. Elucidation of the Phenotypic, Functional, and Molecular Topography of a Myeloerythroid Progenitor Cell Hierarchy. Cell Stem Cell 1, 428–442 (2007).

17. Hao, Y. et al. Integrated analysis of multimodal single-cell data. Cell 184, 3573–3587.e29 (2021).

18. Barbosa, C. M. V. et al. Extracellular annexin-A1 promotes myeloid/granulocytic differentiation of hematopoietic stem/progenitor cells via the Ca2+/MAPK signalling transduction pathway. Cell Death Discov. 5, 135 (2019).

19. Olsson, A. et al. Single-cell analysis of mixed-lineage states leading to a binary cell fate choice. Nature 537, 698–702 (2016).

20. Kwok, I. et al. Combinatorial Single-Cell Analyses of Granulocyte-Monocyte Progenitor Heterogeneity Reveals an Early Uni-potent Neutrophil Progenitor. Immunity 53, 303–318.e5 (2020).

21. Palii, C. G. et al. Single-Cell Proteomics Reveal that Quantitative Changes in Co-expressed Lineage-Specific Transcription Factors Determine Cell Fate. Cell Stem Cell 24, 812–820.e5 (2019).

22. Velten, L. et al. Human haematopoietic stem cell lineage commitment is a continuous process. Nat. Cell Biol. 19, 271–281 (2017).

23. Malinge, S. et al. Ikaros inhibits megakaryopoiesis through functional interaction with GATA-1 and NOTCH signaling. Blood 121, 2440–2451 (2013).

24. Derks, J. et al. Increasing the throughput of sensitive proteomics by plexDIA. Nat. Biotechnol. 41, 50–59 (2023).

25. Thielert, M. et al. Robust dimethyl-based multiplex-DIA doubles single-cell proteome depth via a reference channel. Mol. Syst. Biol. 19, MSB202211503 (2023).

26. Lambert, S. A. et al. The Human Transcription Factors. Cell 172, 650–665 (2018).

27. MacLean, B., et al. Skyline: An open source document editor for creating and analyzing targeted proteomics experiments. Bioinformatics 26, 966–968 (2010).

28. Frankenfield, A. M., Ni, J., Ahmed, M. & Hao, L. Protein Contaminants Matter: Building Universal Protein Contaminant Libraries for DDA and DIA Proteomics. J. Proteome Res. 21, 2104–2113 (2022).

